# A vaccine for global eradication of TB - A novel conceptual framework and design of a potent peptide-based vaccine with universal coverage through advanced computational vaccinology

**DOI:** 10.64898/2026.05.03.722563

**Authors:** Pooja Pawar, Sandhya Samarasinghe

## Abstract

Tuberculosis (TB) remains a formidable global health challenge, exacerbated by the emergence of drug-resistant *Mycobacterium tuberculosis* strains that threaten to render existing drug therapies and vaccine ineffective. Despite the availability of the Bacillus Calmette-Guérin (BCG) vaccine, its limited efficacy—primarily in infants and young children—falls short of reducing TB prevalence or offering adequate protection to adults. Therefore, developing a new TB vaccine with enhanced efficacy and the capability to generate a robust reservoir of memory cells is essential. Addressing the challenge of drug-resistant tuberculosis requires a deep understanding of bacterial evolution and developing robust countermeasures. This study aims to design a next-generation TB vaccine that provides broad-spectrum protection against various *Mycobacterium tuberculosis* strains, including drug-resistant ones. By conducting an in-depth investigation into pathogen-human interactions, the research proposes a holistic framework that leverages computational vaccinology to tackle challenges posed by pathogen polymorphism and overcome the limitations of conventional vaccines. By targeting conserved proteins across diverse TB strains and enhancing both humoral and cell-mediated immunity, this study proposes a new strategy for an epitope-based vaccine that provides long-lasting, universal coverage. An extensive proteomic, reverse vaccinology and immunoinformatics analysis of 159 TB strains yielded 27 highly conserved, immunogenic, non-toxic, and non-allergenic epitopes. These epitopes, consisting of 14-cytotoxic T-lymphocytes (CTL), 5-helper T-lymphocytes (HTL), and 8-B-cell epitopes, were used to construct a three-dimensional, multi-epitope TB vaccine designed based on a new concept introduced in this research for maximising vaccine efficacy. Molecular docking and immune simulation studies demonstrated a significant affinity between the vaccine constructs and toll-like receptors, indicating a strong potential for effective immune system engagement. The crucial features of the epitope-based TB vaccine constructed in this research include sequence conservancy, robust antigenicity, exclusion of self-peptides and potential for diverse allelic interactions. The proposed epitope-based vaccine is poised to be highly effective, safe, and capable of providing universal coverage, potentially paving the way for global TB eradication. Validation in laboratory and clinical settings will be essential to confirm its efficacy and real-world applicability.

## 1. Introduction

Tuberculosis (TB) is an infection of the lungs caused by *Mycobacterium tuberculosis.* As of 2026, TB is recognized as the world’s deadliest infectious disease. Regardless of the advancements made in medical sciences, TB remains the cause of death of more than 1.4 million people annually [1]. Following a temporary rise in COVID-19 deaths, TB has resurged as the top infectious disease killer in 2025, and it remains the foremost cause of mortality in low- and middle-income countries and poses a significant worldwide health concern [1]. Today, vaccine and drug therapy are the two most critical human countermeasures against TB. However, *Mycobacterium tuberculosis* has developed a number of ways to destabilise the human immune response, adapt to the host’s changing environment and disseminate the infection throughout the body. This high level of adaptation has led to the emergence of drug resistant strains of *Mycobacterium tuberculosis* (*Mtb)*.

The widespread emergence of drug-resistant strains has challenged the view of tuberculosis as a treatable disease. Urgency is great for novel solutions but there must be a shift in perspective on this old problem to find effective new solutions. Specifically, a fundamental approach that counters bacterial adaptation at its root is needed to develop an effective and safer vaccine for global and permanent eradication of TB. A vaccine that strikes a blow for immune enhancement against bacterial survivability is key to success in this endeavour. Accordingly, generating a robust and swift immune response and preventing disease progression and transmission are a key consideration for a TB vaccine. Therefore, developing a new TB vaccine with improved efficacy in creating a pool of memory cells is critical. This study primarily aims to identify such potential vaccine candidates against *Mycobacterium tuberculosis* with additional objectives of minimising vaccine targets, identifying protective antigens that can be safely expressed in the laboratory and reducing experimental tests and associated costs of validating vaccine targets. With the ultimate goal of significantly reducing TB globally or drastically minimizing its prevalence, or even potentially eradicating TB altogether, the vaccine development process in this study is focused on pathogen polymorphism, broad coverage of the host population and enhancing humoral and cell-mediated immunity. To achieve this, an epitope-based vaccine that can deliver the intended blow for immune enhancement against bacterial survivability was developed using computational vaccinology.

TB is a highly contagious disease that spreads through the air when an infected person releases airborne droplet nuclei. When an uninfected individual inhales these droplets, they travel down the person’s trachea to the lungs, where TB bacteria multiply, triggering the body’s immune response. Macrophages surround the bacteria, forming a granuloma to contain the spread of TB infection in the lungs [2]. In the early 19^th^ century, TB was a widespread infection. First-line TB drugs for treating drug-susceptible TB patients are isoniazid (INH), ethambutol (EMB), pyrazinamide (PZA) and rifampicin (RIF) [3]. However, the evolution of TB bacteria has led to the emergence of drug-resistant TB strains, rendering these drugs ineffective. Bacteria achieve this through various mechanisms, including making the cell wall impermeable, preventing drug entry into the cells, mutations in the drug target protein, inactivation of drug molecules with the help of bacterial enzymes, and using a transmembrane drug efflux system to expel the drug out from the bacterial cell. In 2025, an 88% treatment success rate was observed in drug-susceptible TB patients, with the remaining 12% attributed to drug resistance and deaths [1].

Vaccines are cost-effective pharmaceutical products essential for the elimination and eradication of infectious diseases. BCG, a hundred-year-old vaccine, has failed to provide complete protection against *Mycobacterium tuberculosis*. The formulation of the live-attenuated BCG vaccine involves the cultivation and attenuation (reducing the virulence) of *Mycobacterium bovis*, the causative agent of TB in cattle [4], using various *in vitro* techniques in the laboratory. BCG has demonstrated its effectiveness by reducing the incidence of tuberculosis in children, miliary TB and tubercular meningitis [5]. However, protection by BCG is highly variable in adults, and it does not significantly prevent chronic infection. The reasons for BCG’s inefficiency in reducing the prevalence and emergence of the disease and protecting adults are currently not well understood. Factors contributing to the high variability in BCG vaccine effectiveness include differences in clinical assays, genetic variability in a sample population, over-attenuation of the currently used BCG strain, different levels of protection against the clinical forms of tuberculosis, variability in *Mycobacterium tuberculosis* strains and malnutrition [6,7].

To accomplish the WHO’s goal of eliminating TB with its End-TB strategy, a vaccine is needed that can stimulate a specific cell-mediated and humoral immune response in the host to eliminate the chances of infection and re-infection from tuberculosis. The traditional vaccine development method requires cultivating pathogens in the laboratory and conducting various biochemical, microbiological and immunological tests to identify vaccine candidates. This approach is laborious, requires expensive experimental testing and sometimes fails to reveal suitable antigens. Currently, about 16 vaccines are in different stages of clinical trials [1]. Among these, seven are protein and viral vector vaccines, while the remaining vaccines are live-attenuated and heat-killed. The main limitation of live-attenuated and inactivated or heat-killed vaccines is the challenge of maintaining purity and safety while culturing the microorganism in the laboratory. A significant drawback of live attenuated vaccines is the reversion of the pathogen into its original pathogenic form [8]. Further, vaccines in clinical trials predominantly rely on two to three antigens for producing recombinant subunits and viral vector vaccines. Therefore, these vaccines do not guarantee broad coverage of immune response. The issue of pathogen polymorphism and drug resistance has also not been addressed adequately by vaccines in clinical trials. Thus, a new and improved vaccine is needed that elicits a more specific immune response and effectively targets a broad spectrum of bacteria.

This study aims to bridge the gaps in vaccine development and proposes a potential solution to reduce the burden of tuberculosis disease globally. The objective is to develop a robust, effective and universal TB vaccine that can bring down the incidence rate of infection, eradicate latent infection, quell the spread of drug-resistance pathogens and prevent the overuse of antibiotics by eliciting a specific and swift immune response while averting autoimmunity or hypersensitive reactions. With this goal in mind, a conceptual and computational framework for designing a universal vaccine against *Mycobacterium tuberculosis* was developed.

In this study, a holistic investigation was first performed to gain deeper insights into the pathogenesis of tuberculosis and the challenges in TB vaccine development. Based on this understanding, a computational vaccinology framework was developed to identify potential vaccine candidates and design a vaccine for TB. The current explosion in bioinformatics has revolutionised the field of vaccine development. Utilising a myriad of bioinformatics tools, the computational vaccinology facilitates the identification of potential vaccine targets without culturing *Mycobacterium tuberculosis* in the laboratory. The main advantage of computational vaccinology is the scope for improving the prediction of vaccine candidates by performing additional computational steps to expand the computational pipeline. Although the proteome of *Mtb* has been considered in vaccine development, its great potential has not been fully utilised in identifying novel vaccine candidates. The proteome includes many proteins contributing to the bacterium’s virulence, progression, and immune evasion. These proteins, such as secretory proteins, lipoproteins, and PE/PPE proteins, represent potential targets for vaccine development. However, their inclusion in vaccine strategies has been limited, and further exploration is necessary to fully understand their potential for developing effective TB vaccines.

The present study utilised computational vaccinology across a wide range of *Mycobacterium tuberculosis* strains to develop an epitope-based vaccine that offers broad-spectrum coverage and enhanced safety. This study stands out as it helps design a TB vaccine that could target multiple highly conserved epitopes critical for inducing both humoral and cell-mediated immune responses. Unlike traditional vaccines that often rely on a limited number of antigens, our approach includes epitopes selected for their immunogenicity, ensuring a robust and long-lasting immune response. Furthermore, our vaccine design incorporates epitopes from proteins essential for the bacterium’s survival and pathogenicity, significantly enhancing its effectiveness against diverse and drug-resistant strains of TB. While epitopes have been considered in some prior vaccine strategies, they have not been fully leveraged to create a comprehensive vaccine with the specificity, safety, and broad-spectrum efficacy achieved in this study. Using advanced computational methods, our vaccine design overcomes previous limitations by systematically identifying and optimizing epitopes that provide universal protection against a wide array of TB strains, including those resistant to current treatments.

This study is designed to address the challenges of conventional vaccine development that include expensive, time-consuming and arduous experimental testing and safety concerns associated with culturing the pathogen in a laboratory. Our study focuses on identifying surface-exposed, secreted, and adhesin proteins from the TB bacterium’s proteome that can help pinpoint conserved epitopes even in highly variable or drug-resistant strains of *Mycobacterium tuberculosis*. While these concepts have been considered, previous vaccine efforts have not fully exploited these strategies due to a limited understanding of host-pathogen interactions. By leveraging our comprehensive analysis of these interactions, we developed a novel approach that addresses the shortcomings of earlier vaccines, thereby enhancing efficacy against diverse and resistant TB strains. Unlike traditional methods, computational vaccinology saves time and costs by minimising repeated laboratory testing. By designing a series of computational steps to address these challenges, this study develops a powerful and safe epitope-based vaccine and creates a compact formulation that further enhances its effectiveness.

Today, after 100 years of research in the field of TB vaccines, designing and developing an ideal universal TB vaccine is still hindered by several challenges, such as genetic diversity of pathogen, variability in sequence conservancy, immunodominance issues, antigenicity, the need to exclude self-peptides and multiple allelic interactions. This study addresses these challenges by employing an in-depth analysis of host-pathogen interactions, revealing distinctive strategies *Mycobacterium tuberculosis* employs to evade immune responses. This understanding, the knowledge of the limitations of BCG, aforementioned challenges in developing an effective vaccine and the advantages of computational techniques in vaccine design, helped us create a conceptual framework for the design of a new TB vaccine with universal coverage.

In this study, some crucial concepts were used for designing an effective and robust TB vaccine compared to the conventional TB vaccine development approach as summarised below:

### (i) Capturing *Mycobacterium tuberculosis* genetic variability

Genetic variability among different strains of the immune evasive *Mycobacterium tuberculosis* increases the chances of drug resistance. Around 0.3 million people die each year from drug-resistant TB infections. High genetic variability is one of the fundamental reasons behind the non-availability and less effective vaccines against TB. The practical solution to the TB bacteria polymorphism issue is to use highly conserved vaccine targets from *Mycobacterium tuberculosis* proteome to develop a universal TB vaccine.

### (ii) Surface-exposed protein as a potential candidate for epitope-based vaccine

Most of the proteins involved in the pathogenesis of TB are present in the cell membranes or cell walls. Surface proteins are easily accessible to the host immune system. For example, the extracellular proteins of *Mtb* are involved in interactions with toll-like receptors (TLR-2 and TLR-4) on the macrophages [9,10,11]. Thus, surface-exposed proteins are potential vaccine targets to weaken *Mtb*.

### (iii) Identification of proteins involved in pathogenesis

The proteome of *Mtb* includes proteins involved in virulence, TB progression, and evasion of the host immune response. Some examples are secretory proteins, lipoproteins, PE (proline-glutamate) and PPE (proline-proline-glutamate) proteins. Secretory signal peptides are ubiquitous protein-sorting signals that help translocate a protein across the cytoplasmic membrane to the cell wall and extracellular space in mycobacteria. Studies have shown that the secretory systems of *Mtb* are involved in the virulence mechanism [12]. *Mtb* has two important secretion systems: the secretory pathway (Sec pathway) and the twin-arginine translocase pathway (TAT pathway). The Sec pathway helps in the translocation of proteins in their unfolded state. The TAT pathway translocates proteins in their folded state. The proteins involved in the TAT pathway help translocate the phospholipase C enzyme in *Mtb*. This enzyme exhibits a cytotoxic effect on the host’s alveolar macrophage [13]. Lipoproteins, a significant set of membrane proteins in *Mtb*, perform various functions, including modulating the inflammatory response, translocating virulent factors in the macrophages, conferring drug resistance, signal transduction, and nutrient uptake [14]. These proteins constitute 2.5% of the predicted proteome of *Mtb* and represent a significant virulent protein family [15]. The PE and PPE protein families in the cell wall and cell membrane [16] contribute to antigen variability in *Mtb* [17].

### (iv) Eliciting a comprehensive immune response

Triggering a robust immune response is the most crucial challenge in vaccine development. The conventional TB vaccine approach focused on the helper T-cell immune response has not shown protective immunity against *Mtb* [18]. Thus, an epitope-based TB vaccine containing helper-T cell, cytotoxic-T cell and B-cell epitopes can stimulate a specific cell-mediated and humoral immune response in the host to eliminate the chances of infection and re-infection of tuberculosis.

### (v) Careful selection of effective vaccine candidates

Major histocompatibility complex (MHC) class-I molecules are present on the surface of all nucleated cells and play a crucial role in the immune system by presenting intracellularly processed antigens to cytotoxic T-Lymphocytes (CTLs). This process typically occurs when a TB bacterium infects a cell. Inside the cell, proteins from bacteria are degraded into smaller peptides, which are then loaded onto MHC-I molecules. These MHC-I molecules, with the bound antigen peptides, are transported to the cell surface, where they are recognized by CTLs. The recognition triggers CTLs to initiate a response that destroys the infected or abnormal cells [19].

MHC-II molecules are present primarily on the surface of antigen-presenting cells (APCs) such as dendritic cells, macrophages, and B cells. They play a crucial role in the immune system by presenting extracellularly derived antigens to Helper T-Lymphocytes (HTLs) This process typically occurs when an APC engulfs and internalises TB bacteria. Inside the APC, antigenic proteins are degraded into smaller peptides known as epitopes, which are the part of an antigen that binds to T-cells, B-cells, or antibodies in the host body -T-cells and B-cells are the principal constituents of the immune system. These epitopes are then loaded onto MHC-II molecules, which are transported to the cell surface to be presented to HTLs. This recognition activates HTLs, which then release cytokines that help regulate and coordinate the immune response, including activating B cells to produce antibodies and enhancing the activity of other immune cells [19]. Therefore, careful selection of the immunodominant T-cell epitopes and determining their binding affinity with MHC molecules is crucial in vaccine design and development.

### (vi) Capturing genetic variability of the human population

Another important concept in developing an effective TB vaccine is broad coverage in the human population. In most cases, a single strain or single antigen vaccine is effective in a small subset of patients in a specific region. Such a vaccine may be considered ethnically biased in terms of protection. MHC molecules are highly polymorphic, with over a thousand human leukocyte antigens (HLA) alleles identified. Different HLA types are expressed at varying rates across different ethnicities worldwide. Therefore, selecting T-cell epitopes binding to MHC HLA alleles most prevalent in the human population is proposed to help design a universal vaccine with optimal efficacy against TB.

### (vii) Selecting safer vaccine candidates to reduce the probability of side effects

An epitope-based vaccine is safer than live-attenuated or heat-killed vaccines as they only contain fragments from antigenic proteins and not the live components of the pathogen. Thus, there is no risk of reversal of an epitope-based vaccine to harmful elements. Additionally, eliminating cross-reactive and toxic epitopes that cause autoimmunity or hypersensitive reactions in the host makes the vaccine safer.

### (viii) Immunogenicity enhancement

The use of adjuvants in vaccines heightens the immune response [20]. An adjuvant increases the vaccine’s immunogenicity and helps induce a robust and long-lasting immune response in immunocompetent individuals.

Using the aforementioned concepts, a systematic computational pipeline, involving comparative proteomics, reverse vaccinology, immunoinformatics and structural vaccinology, was designed to develop an epitope-based TB vaccine from the proteome of *Mycobacterium tuberculosis*. The proposed method involves six phases (Figure 1). The first phase involved comparative proteomic analysis in identifying conserved proteins across 159 *Mycobacterium strains* and functional categorisation of these conserved proteins. Targeting highly conserved regions for their significant structural and functional roles in the pathogen’s life cycle would provide a broad spectrum of protection against many *Mycobacterium tuberculosis* strains and drug resistance. In the second phase, reverse vaccinology was used on conserved proteins to analyse the surface-exposed, antigenic, non-allergic proteins involved in the pathogenesis of TB that were easily accessible to surveillance by the host immune system. A vaccine developed based on many strains with broad coverage and a deep understanding of host-pathogen interaction can significantly contribute to eliminating TB globally or drastically minimise its prevalence.

**Figure 1:**
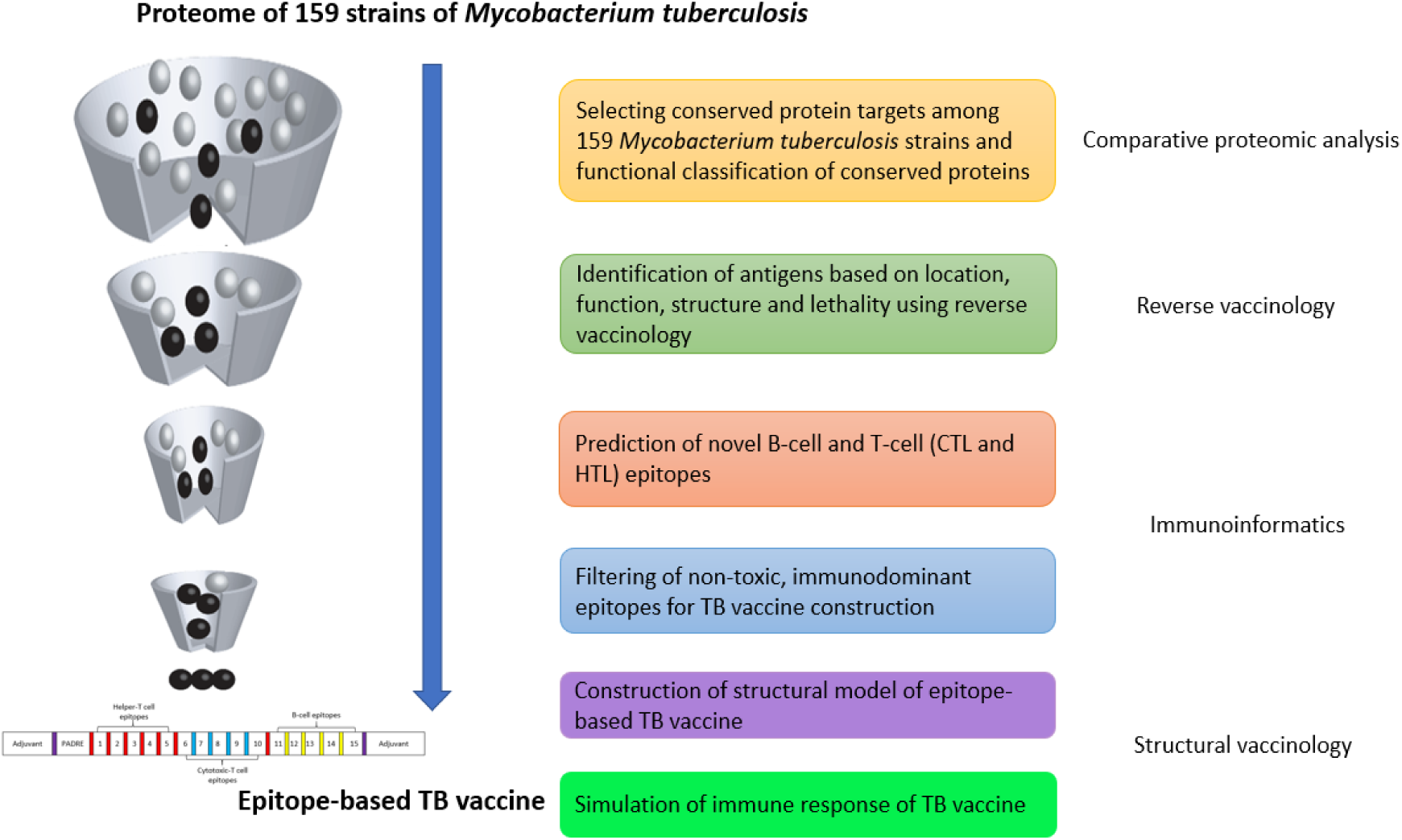
Conceptual framework for constructing an epitope-based TB vaccine against *Mycobacterium tuberculosis* using comparative proteomic analysis, reverse vaccinology, immunoinformatics and structural vaccinology

An immunoinformatic approach was employed in phase three to predict B-cell and T-cell epitopes. T-cell epitope selection to a broad range of MHC HLA alleles is crucial for global coverage of the human population. The fourth phase comprised filtering the immunodominant epitopes that can induce protective and robust immunity in the host. The B-cell and T-cell epitopes predicted in this phase were used to generate the final vaccine construct. In the fifth phase, using structural vaccinology, the structural model of the new TB vaccine, with the adjuvant attached to it, is generated. The binding affinity of the TB vaccine with toll-like receptors in the host antigen-presenting cell helps determine the behaviour of the vaccine inside the host body. The sixth phase involved analysing the immune response generated by the designed epitope-based TB vaccine using the C-ImmSim server. This step determines the efficacy of the developed TB vaccine inside the human body. Details of the methodology and bioinformatics tools used in each step can be found in the next section.

## 2. Materials and Methodology

The methodological framework shown in Figure 1 was employed to identify potential T-cell and B-cell epitopes for developing a universal vaccine against tuberculosis. This approach utilised several bioinformatics tools to analyse the proteome of 159 strains of *Mycobacterium tuberculosis*, thereby addressing the challenges of antigen variability, drug resistance and limited coverage. The detailed steps of the methodology presented in Figure 1 are elaborated below.

### 2.1 Comparative proteomic analysis

The proteomes of the completely sequenced 159 strains of *Mycobacterium tuberculosis* were retrieved in FASTA format via the NCBI (National Center for Biotechnology Information Genome FTP site). Standalone BLAST (Basic Local Alignment Search Tool) was used to perform a homology search among *Mycobacterium tuberculosis* strains. *Mycobacterium tuberculosis* H37Rv served as a reference proteome for analysis. Protein sequences with a sequence similarity greater than 99% across all 159 strains were considered conserved proteins and selected for further research. The resulting conserved proteins were categorised into eleven distinct functional categories given by Smith (2003): lipid metabolism; information pathways; cell wall and cell processes; stable RNAs; IS elements and bacteriophages; PE and PPE proteins; intermediary metabolism and respiration; regulatory proteins; virulence, detoxification and adaptation; conserved hypothetical function; and proteins of unknown function. The categorisation was done based on the sequence similarity results obtained through BLAST.

Hypothetical proteins (HPs) and proteins with unknown functions were particularly interesting, as they may play crucial roles in essential metabolic activity, growth and development, or pathogenicity and could contribute to improving the understanding of the survival of TB bacteria. For this, function annotation was performed using domain prediction tools such as Pfam [21], Interpro Scan [22] and SMART [23], combined with homology search through BLAST [24].

### 2.2 Identifying antigenic proteins using reverse vaccinology

A reverse vaccinology approach was applied to identify outer-membrane, antigenic and non-allergenic proteins possessing unique characteristics stated below following a systematic approach as presented in this section.

- **membrane-spanning regions-** these regions allow the protein to anchor in the membrane, making them prime targets for immune recognition since they are exposed to the host immune system
- **signal peptides-** essential for proteins that are presented on the cell surface or secreted, making them accessible to the immune system
- **lipoprotein signatures-** play significant roles in *Mycobacterium tuberculosis*-host interactions and are often immunogenic, making them suitable targets for TB vaccine development
- **adhesion probability-** targeting proteins with high adhesion probability, such as adhesins, in a vaccine can block the TB bacteria’s attachment and invasion of host cells, thereby preventing infection and enhancing vaccine effectiveness.

#### 2.2.1 Subcellular localisation (SCL) prediction

Cell membrane and extracellular space proteins are essential to maintaining cell integrity. They are considered suitable TB vaccine candidates since they are easily accessible to the host’s immune system. SCL predictions were made using PSORTb v.3.0 [25], CELLO [26], LocTree3 [27], SOSUI [28], pLoc_bal-mGpos [29] and GramLocEN [30]. PSORTb v.3.0. CELLO and LocTree3 employ machine learning models (support vector machines (SVM)) for SCL prediction, while SOSUI and pLoc_bal-mGpos rely on protein sequence information, such as physiochemical properties, for subcellular localisation prediction. GramLocEN uses a single and multi-label elastic net classifier to make SCL predictions. Proteins predicted to be located in the cell membranes and extracellular spaces by at least four tools were selected for further analysis.

#### 2.2.2 Transmembrane (TM) helix prediction

Transmembrane proteins containing a single transmembrane α-helix were prioritized, as proteins with multiple transmembrane helices pose challenges for laboratory cloning and purification [31]. Thus, selected outer membrane and extracellular proteins having a single transmembrane α-helix were used to help design the vaccine. TMHMM, based on the hidden Markov model (HMM), was used to predict the transmembrane regions and their orientation [32].

#### 2.2.3 Identification of functionally relevant proteins

*Mycobacterium tuberculosis* outer membrane and extracellular space consist of some essential signal peptides, secretory proteins involved in Sec and TAT pathway and lipoproteins. The functionally important proteins were identified using: (i) SignalP 4.1 server which is a combination of several artificial neural networks (ANN) and predicts the presence and position of a signal peptide in the query sequence [33]; (ii) PRED-TAT predicts TAT-signal peptides using HMM [34]; (iii) SecretomeP is an ANN-based method for predicting secretory proteins [35], and proteins with a score of 0.5 or above were selected; and (iv) PRED-LIPO is an HMM-based method for predicting lipoproteins [36].

#### 2.2.4 Antigenicity and allergenicity prediction

To develop a TB vaccine capable of eliciting a specific immune response, it was necessary to identify antigenic proteins with infection-inducing potential. Three bioinformatics tools were used to predict antigenic proteins: VaxiJen helps predict the antigenic properties based on the physiochemical properties of the protein [37]; VirulentPred is an SVM-based method [38], whereas the MP3 tool uses SVM and HMM [39] to predict antigenicity within a protein.

Proteins predicted as antigenic by all three tools underwent allergenicity prediction using Algpred, which utilizes a machine learning model (random forest) to differentiate between allergenic and non-allergenic protein based on the amino acid composition (AAC) of the protein [40].

#### 2.2.5 Adhesion probability

Mycobacteria have adhesin proteins that help them attach to the host cell-surface receptors. The adherence of *Mycobacterium tuberculosis* to host cells generates membrane agitation and enables a robust interaction between *Mycobacterium tuberculosis* and the host [41]. The adhesion probability of the selected antigenic and non-allergenic proteins was predicted using an artificial neural network-based method known as SPAAN (software for predicting adhesins and adhesin-like proteins using neural networks) [42]. The proteins with a probability score of 0.5 or above were selected for further study.

#### 2.2.6 Interaction pathway analysis

Proteins with more than ten interacting partners and a confidence score of 0.4 or above were chosen. Proteins interacting with many partners are often more central to the biological network they are involved in, playing critical roles in essential processes and maintaining the pathogen’s survival. Therefore, understanding these interactions can provide insights into the functional significance of the selected antigenic and non-allergenic proteins with high adhesion probability in *Mycobacterium tuberculosis*. The protein-interaction network was analysed using the STRING database [43], selecting proteins with more than ten interacting partners and a confidence score of 0.4 or above to identify those likely critical for the pathogen’s function and virulence.

#### 2.2.7 Homology to humans

It is crucial to exclude proteins similar to human proteins in function or involved in common metabolic pathways to avoid triggering autoimmunity or hypersensitive reactions in the host. To achieve this, BLASTp was used to identify sequence similarities between *Mycobacterium tuberculosis* proteins and human proteins; proteins with more than 30% sequence similarity were eliminated from the study. The Kyoto Encyclopaedia of Genes and Genomes (KEGG) [44] was used to identify and exclude proteins involved in metabolic pathways common to humans and *Mycobacterium tuberculosis*. At the end of the reverse vaccinology step, the remaining set of antigens includes those that are located on the plasma membrane, have a single transmembrane α-helix, are functionally necessary to the bacterium, have a high adhesion probability to host receptors and do not pose a risk of causing adverse reactions in the host.

### 2.3 Prediction of B-cell and T-cell epitopes and filtering using Immuno-informatics

Antigens may contain epitopes that invoke humoral immune response (B-cell epitopes) and cell-mediated immune response (T-cell epitopes). Therefore, antigens identified using the reverse vaccinology approach were further analysed to predict and filter potential B-cell and T-cell epitopes using Immunoinformatics tools to develop an effective vaccine against TB.

#### 2.3.1 B-cell epitope prediction

B-cell epitopes play a vital role in initiating the humoral immune response when they are bound to B-cells of the host immune system. Three tools were used to predict B-Cell epitopes: The Bepipred tool of the Immune Epitope Database (IEDB) uses a hidden Markov model and a propensity scale method to predict linear B-cell epitopes accurately [45]; ABCpred uses a recurrent neural network for predicting B-cell epitopes [46]; and BCPREDS is a server used for B-cell epitope prediction using the amino acid pair (AAP) antigenicity scale [47]. The common epitopes predicted by all three methods were chosen.

#### 2.3.2 T-cell epitope prediction

T-cell epitopes initiate a cell-mediated immune response. For this to happen, they must bind to MHC molecules, an essential class of proteins present on the surface of a cell and play a crucial role in cell-mediated immunity. The vital function of the MHC molecules is the presentation of fragmented or processed antigens to the appropriate T-cells of the immune system. The binding of a peptide with an MHC molecule is necessary for recognising an epitope by the T-cell receptors. MHC-I and MHC-II are two types of molecules that recognise T-cell epitopes and therefore, epitopes in the selected antigens that bind with these two molecular types must be identified.

### **(i)** MHC-I binding T-cell (cytotoxic-T cell) epitope prediction

MHC-I molecules are present on the surface of all nucleated cells in the human body. MHC-I molecules present T-cell epitopes to the cytotoxic T-lymphocyte (CTL). The triggering of CTL causes the antigen-presenting cells (APCs) to undergo programmed cell death. Three methods were used in this step to identify MHC-I binding (cytotoxic) T-cell epitopes in the selected antigens: IEDB MHC-I [48], NetCTLpan 1.1 [49] and IEDB MHC-NP [50]. In IEDB MHC-I, a reference file containing 27 MHC-I HLA alleles was used for a broader human population coverage. The 9-mer epitopes with percentile rank lower than 0.5 were chosen. The lower percentile rank value indicates a higher affinity of the epitope towards the MHC-I molecule. NetCTLpan 1.1 uses an artificial neural network for predicting MHC-I epitopes. IEDB MHC-NP is a machine learning method that predicts naturally processed peptides using MHC-I experimental data [50]. Common epitopes across all three prediction results were selected as T-cell MHC-I epitopes.

### **(ii)** MHC-II binding T-cell (helper-T cell) epitope prediction

MHC-II molecules are present on the surfaces of APCs. The primary function of MHC-II is to present the antigen to the naïve T-helper cells. This interaction leads to the release of cytokines that help transform naïve-T-helper cells into effector T-cells or memory T-cells. The NetMHCIIpan 3.2 server supports the prediction of epitopes that bind to MHC-II molecules. The method uses a feed-forward artificial neural network that predicts all three MHC-II alleles HLA-DR, -DP, -DQ, for humans covering 36 HLA-DR, 9 HLA-DP, and 27 HLA-DQ molecules [51]. The second method used for prediction was the IEDB MHC-II binding server for predicting 15-mer MHC-II epitopes using the consensus prediction method [52]. Thirty MHC-II HLA alleles from the reference set were used in our study. The epitopes with percentile rank lower than 0.5 were chosen. The lower percentile rank value indicates a higher affinity of the epitope towards the MHC-II molecule. The epitopes commonly predicted by both methods were selected as MHC-II epitopes.

#### 2.3.3 Filtering of epitopes

Although antigens were initially selected based on specific properties, individual epitopes must be further evaluated for antigenicity, non-toxicity, and non-allergenicity as well as hydrophilicity and population coverage to ensure their suitability for vaccine development. The filtering process had the following steps:

##### 2.3.3.1 Antigenicity and toxicity prediction of B-cell and T-cell epitopes

VaxiJen [37] was used to predict the epitopes having the potential to initiate an immune response (antigenicity). The epitopes with a VaxiJen score of 0.8 or above were selected. The toxicity and non-toxicity of an epitope can be determined using the SVM-based tool, ToxinPred [53].

##### 2.3.3.2 Hydrophilicity prediction

Hydrophilic epitopes interact well with water molecules, which usually indicates that they are located on the surface of proteins. This surface exposure makes them more accessible to the immune system, making them ideal candidates for vaccine development. The grand average hydropathicity (GRAVY) index score [54] was predicted using ProtParam [55].

##### 2.3.3.3 Allergenicity prediction

Allergenicity determines whether the epitope is allergic or non-allergic to the host. To make sure that the designed TB vaccine does not elicit an allergenic response inside the host body, AllerTOP 2.0 was used. AllerTOP 2.0 predicts epitope allergenicity based on the query sequence’s physiochemical properties [56].

##### 2.3.3.4 Population coverage analysis

Designing a universal vaccine for TB needs an analysis of the population coverage to ensure broad spectrum coverage and minimise the risk of developing an ethnically biased vaccine [57]. The population coverage assessment used the IEDB epitope analysis resource for a distinct population coverage. This analysis tool predicts the population coverage, average number of epitope hits or HLA combinations recognised by diverse ethnic groups or populations, and the least number of epitope hits expected for 90% of population coverage [57]. The final epitopes for a vaccine must have extensive population coverage worldwide.

### 2.4 *In silico* vaccine construction using structural vaccinology

After selecting epitopes from the Immunoinformatics analysis, structural vaccinology was implemented to design the structural model of the TB vaccine and analyse its interaction with toll-like receptors present in the host. Further, codon optimisation was conducted to ensure optimal gene expression of the vaccine inside the host body and *in silico* cloning helps design and validate the insertion of the optimised DNA sequence into a vector before actual laboratory work, saving time and resources.

#### 2.4.1 Vaccine structure construction and validation

##### 2.4.1.1 Vaccine design

For designing a final vaccine sequence from the shortlisted epitopes, a new approach with the following steps was developed:

i. *Selection of linkers*: Flexible linkers, such as GPGPG and AAY, were used to connect different types of epitopes within the vaccine construct. These linkers help maintain the proper folding and function of the protein domains, ensuring that the epitopes remain accessible for immune recognition. Figure 2 shows the linker used for designing the vaccine proteins. The GPGPG linker was used to attach helper-T cell epitopes with CTL epitopes and CTL epitopes with B-cell epitopes. The GPGPG linker was also used to connect helper-T cell epitopes with themselves, and the AAY linker was used to connect CTL epitopes to themselves. For B-cell epitopes, a KK linker was used. The 3D structure of each epitope with the attached linker was generated using PEPstrMOD [58].
ii. *Pan DR epitope (PADRE) attachment:* PADRE is a universal synthetic 13 amino acid peptide. The PADRE sequence was used to improve the efficacy of cell-mediated immune response. The attachment of PADRE with HTL epitopes would facilitate quick and robust interaction with Toll-like receptors in the host body. It would accomplish the goal of initiating the immune response. Thus, it was attached to the first HTL epitope in the vaccine protein [59].
iii. *Initialising the vaccine sequence:* For this, the 3D structure of the PADRE sequence with the GPGPG linker attached to its end for connecting with the first helper-T cell epitope was generated using PEPstrMOD. Next, molecular docking of the 3D structure of PADRE was completed with the structure of each helper-T cell epitope (structure constructed in step (i)) using PatchDock [60] and FireDock [61]. The helper-T cell epitope with strong binding with the structure of PADRE was selected as the first epitope in the vaccine sequence design.
iv. *Finalising the vaccine sequence:* After the above initialising phase, the PADRE+1st helper-T cell epitope structure was constructed using I-TASSER [62], and its compatibility was rechecked with the remaining helper-T cell epitopes. This process was completed first for helper-T cell epitopes, then cytotoxic-T cell epitopes and finally the B-cell epitopes. This process continued until the last B-cell epitope was found for the vaccine sequence.
v. *Adjuvants for enhancing immune response*: Adjuvants were used to enhance the immune response in the host. In this study, two adjuvants were used. At the N-terminal of the protein, the 50S ribosomal protein L7/L12, and at the C-terminal of the vaccine, protein β-defensin [63] was used (Figure 2). 50S ribosomal protein L7/L12 was chosen for its role in enhancing the immune response as an adjuvant, while β-defensin, known for its antimicrobial properties, was selected to boost the vaccine’s efficacy by acting as an additional adjuvant at the C-terminal. The 50S ribosomal protein L7/L12 and β-defensin were attached to the vaccine protein obtained above using an EAAAK linker (purple coloured box in Figure 2).

**Figure 2:**
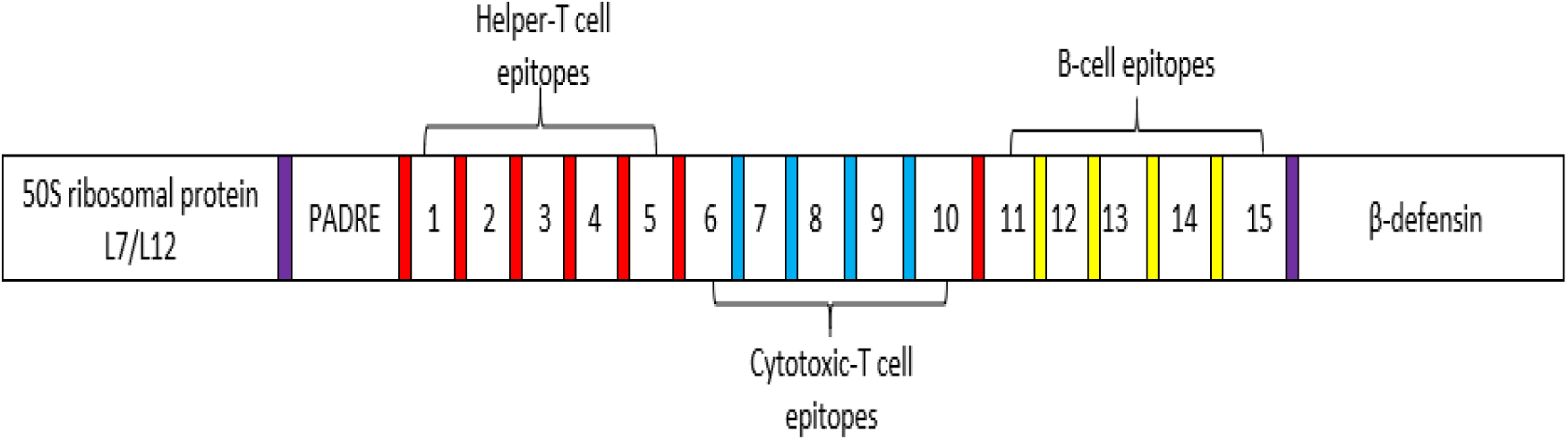
The location of different linkers used in designing the final vaccine protein - purple coloured box: EAAAK linker, red: GPGPG, blue: AAY, yellow: KK. The adjuvants are connected to the vaccine protein using EAAAK. The GPGPG linker combined different types of epitopes (attachment of helper-T cell epitopes with themselves, helper-T cell epitopes with cytotoxic-T cell and cytotoxic-T cell epitopes with B-cell epitopes. Cytotoxic-T cell epitopes were attached using the AAY linker and KK linker was used for connecting B-cell epitopes.

An effective vaccine should possess the following attributes: high antigenicity, non-toxicity, non-allergenicity and an appropriate molecular structure to ensure robust immune responses. The designed vaccine was tested for these attributes.

##### 2.4.1.2 Antigenicity and allergenicity prediction

VaxiJen [37] was used to predict the antigenicity of the final vaccine construct. Algpred, an SVM module, was used to determine the vaccine protein allergenicity [40].

##### 2.4.1.3 Physiochemical properties analysis

ProtParam was used for computing the various physiochemical properties of the final TB vaccine. The properties include molecular weight (MW), the isoelectric point of a protein (pI), amino acid composition, extinction coefficient [64], instability index (II), estimated half-life [65], aliphatic index [66] and the GRAVY index score [54].

##### 2.4.1.4 Secondary structure prediction

The secondary structure provides insights into how the protein folds and maintains stability. In vaccine design, it is crucial to ensure that the epitopes maintain a structure recognisable by the immune system, which often relies on specific structural motifs. The secondary structure prediction of the vaccine construct used PSIPRED [67]. Knowledge of alpha-helix, beta-sheet and coils improves understanding of the structure of the vaccine construct.

##### 2.4.1.5 Tertiary structure construction and validation

The tertiary structure of the final TB vaccine construct was generated using RaptorX [68]. First, RaptorX performed a template search based on the similarity of the input sequence and then constructed a good-quality structural model. The output structural model of the vaccine given by RaptorX was further refined using GalaxyRefine to minimise any distortions present in the structure [69]. The GalaxyRefine server employs molecular dynamic simulation for structural relaxation and changes amino acids with high probability rotamers to refine the three-dimensional structures. The output of GalaxyRefine generally contains five refined models. Other parameters evaluated after refinement are GDT-HA (Global Distance Test-High Accuracy) [83], MolProbidity, clash score, RMSD, poor rotamers, and Rama favoured scores. The Global Distance Test (GDT) evaluates the accuracy of the predicted protein structure by comparing it to a known reference structure, providing a quantitative measure of how closely the predicted model matches the actual structure [84]. MolProbidity measures the quality of a protein structure, particularly the geometry of bond angles and lengths [85]. Clash Score: Indicates the number of steric clashes in the structure where atoms are positioned too closely together. Poor Rotamers refer to side-chain conformations that are unlikely or energetically unfavourable. Rama Favoured Scores, derived from the Ramachandran plot, indicate the percentage of amino acid residues in favourable conformations within the protein structure. RMSD (Root Mean Square Deviation) measures the average distance between the atoms of superimposed proteins, indicating the deviation of the predicted structure from the reference.

In the last step, RAMPAGE was used to determine the quality of the vaccine structural models [70]. If a vaccine structural model had more than 90% of its residues in the most favoured region, it was considered the best quality model and was used for further analysis.

#### 2.4.2 Vaccine-Toll Like receptor (TLR) docking and dynamics

*Mycobacterium tuberculosis* usually interacts with TLR-2, 4 and 6 [10,11]. TLRs on the surface of the antigen-presenting cells (APCs) interact with the pathogen to initiate the innate immune response. The interaction of TLRs with the final vaccine construct was analysed by molecular docking. For the docking process, the 3D structures of TLR-2, TLR-4 and TLR-6 were retrieved from PDB with PDB ID 2Z7X, 4G8A and 4OM7, respectively. The Cluspro 2.0 [71] webserver was used for performing molecular docking of the vaccine construct-TLR. Cluspro 2.0 performs three steps: (i) performing rigid docking using the fast Fourier transform (FFT) correlation method, (ii) calculation of the root-mean-square deviation (RMSD) for docked clusters, and (iii) refinement of the structure using a Monte Carlo simulation algorithm [71]. This refinement step adjusts the orientation and conformation of the vaccine construct and TLR to improve binding accuracy and optimise interactions. PDBsum was used to get schematic illustrations of the docking between TLRs and TB vaccine construct [72]. After docking, normal mode analysis (NMA) was used to evaluate the flexibility and stability of the vaccine structure. Ideally, we expect low mobility and minimal deformation, indicating a stable and robust vaccine candidate. iMODS [73] was used to determine the dynamic motion of the vaccine-TLR docked complex.

#### 2.4.3 Codon optimisation and *in silico* cloning

By optimising the codon usage in the vaccine construct to match the host organism’s preference, the efficiency of protein expression is significantly improved, leading to higher yields of the vaccine protein. To gain insight into the expression level of the vaccine gene inside the host, it is necessary to determine the codon adaptation index of the TB vaccine. Java Codon Adaptation Tool (JCat) performed reverse translation and codon optimisation in *E. coli* strain K12. *E. coli* strain K12 is a well-characterised reference strain commonly used in research and biotechnology. JCat calculates the codon adaptation index (CAI) value and guanine-cytosine (GC) content of the submitted protein sequence [74]. For favourable transcriptional and translational efficiencies, the ideal GC content should be between 30 and 70%, and the CAI score should be higher than 0.8.

Further, *in-silico* cloning was performed using the SnapGene tool. The DNA sequence was inserted into the pET-28a (+) vector, commonly used to express recombinant proteins in *E. coli*. This step validates the potential of the vaccine protein sequence as a promising vaccine candidate through various experimental tests, ensuring it can be produced at a scale suitable for further development.

### 2.5 Analysis of the evoked immune response by the TB vaccine construct

The most crucial step in the study was understanding the immunogenic nature of the final vaccine construct. The C-ImmSim [75] server was used to estimate the constructed TB vaccine’s immunogenic potential. C-ImmSim server is an agent-based model that uses the position-specific scoring matrix (PSSM) and machine learning approaches, such as neural networks, for predicting epitopes, the immune interactions of epitopes and immune cells, and immune response activation [75]. The C-ImmSim server simulated three sections that corresponded to the three different anatomical regions inside the host body: (i) bone marrow, (ii) thymus, and (iii) lymph nodes [75]. For determining vaccine efficacy, the TB vaccine protein sequence in FASTA format was given as input to the server. The vaccine protein sequence was administered three times at an interval of four weeks. In C-ImmSim, one simulation step represented eight hours. So, over a year, 1050 simulation steps were used to predict the immune response to the TB vaccine.

## 3. Results and Discussion

This section presents the results for the new vaccine obtained from the vaccine development pipeline incorporating a range of bioinformatics tools.

### 3.1 Comparative proteomic analysis

Comparative proteomic analysis helps identify conserved vaccine candidates that are broadly effective on different *Mycobacterium tuberculosis* strains and provide protection against drug resistance. The complete proteome sequence of 159 different *Mycobacterium tuberculosis* strains was downloaded from the NCBI Genome FTP site (Supplementary Table S1 provides information on the 159 *Mycobacterium tuberculosis* strains). *Mycobacterium tuberculosis* H37Rv (GenBank accession number NC_000962.3), with 3906 proteins, was taken as the reference proteome for identifying the conserved proteins within the 159 strains. Proteome comparison was made using a standalone BLAST, and the results from sequence similarity were stored in Excel files. Out of 3906 proteins of H37Rv, 1982 (≈ 51%) proteins were conserved among the 159 strains of *Mycobacterium tuberculosis* (Table 1) (Full details are in Supplementary Table S2). After performing sequence similarity among the 159 strains, each protein was categorised into a functional class, as given by Smith (2003). The study aimed to understand the functional significance of conserved proteins and predict their role in the survival strategies of bacteria, particularly how these proteins may contribute to the evolution of different bacterial strains. Table 1 shows the distribution of conserved proteins in each functional category. A high percentage (73%) of conserved proteins were found to be involved in critical functional classes when the conserved proteins (1982) were compared with the whole proteome (3906); for example, intermediary metabolism and respiration (28%), cell wall processes (19%), information pathways (8%), lipid metabolism and regulatory proteins (7%), and virulence, detoxification and adaptation (4%) (as highlighted in Table 1). Comparative proteomic analysis indicates that *Mycobacterium tuberculosis* tends to alter proteins less crucial for the bacteria’s growth and development. *Mycobacterium tuberculosis* safeguards the proteins involved in the normal functioning of bacteria, progression of TB disease and survival within the host.

**Table 1:**
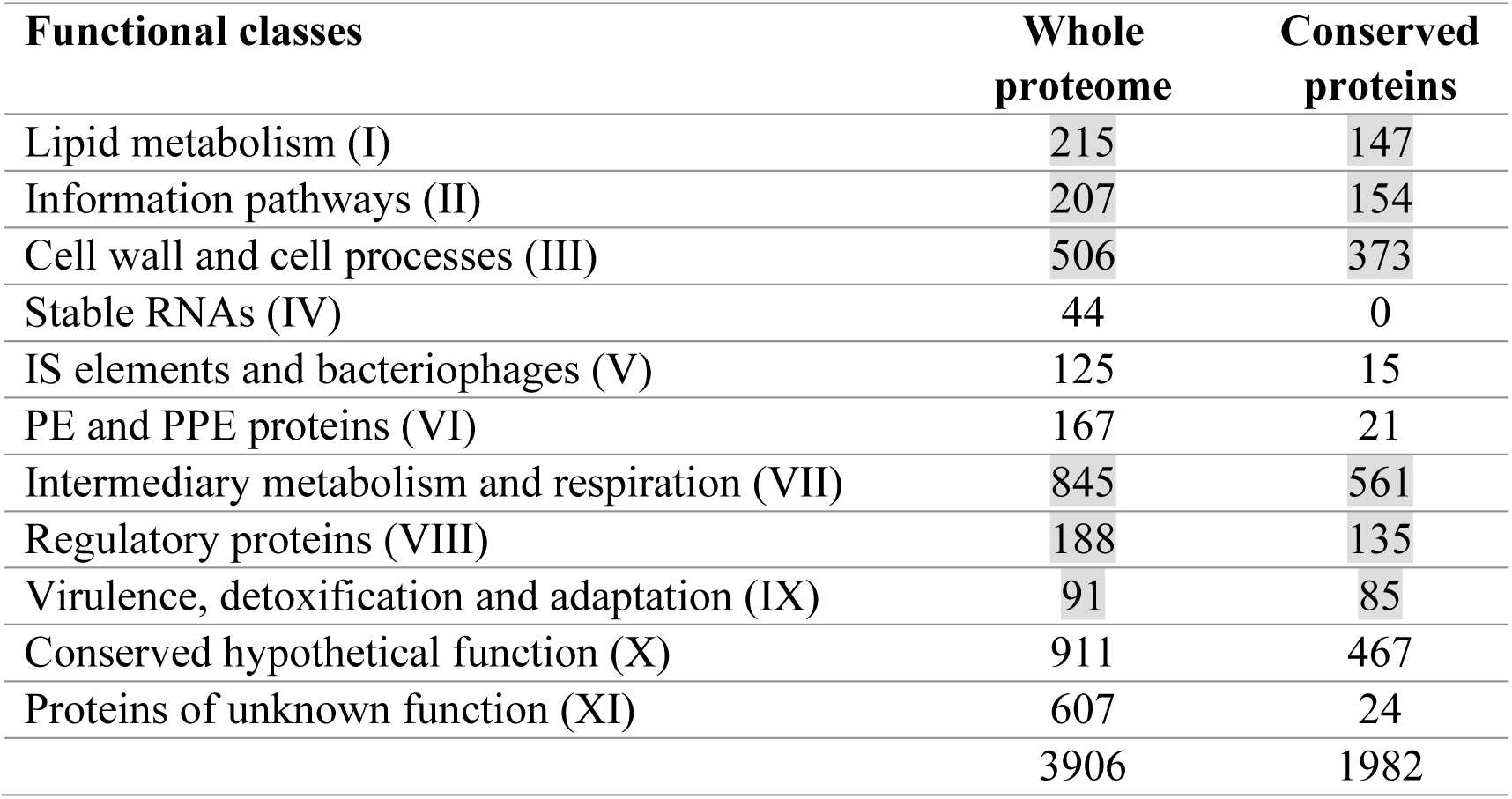
Distribution of whole proteome (3906 proteins) of *Mycobacterium tuberculosis* H37Rv and the conserved proteins (1982 proteins) identified after performing sequence similarity within 159 strains and distributing into functional categories suggested by Smith (2003)

| Functional classes | Whole proteome | Conserved proteins |
| --- | --- | --- |
| Lipid metabolism (I) | 215 | 147 |
| Information pathways (II) | 207 | 154 |
| Cell wall and cell processes (III) | 506 | 373 |
| Stable RNAs (IV) | 44 | 0 |
| IS elements and bacteriophages (V) | 125 | 15 |
| PE and PPE proteins (VI) | 167 | 21 |
| Intermediary metabolism and respiration (VII) | 845 | 561 |
| Regulatory proteins (VIII) | 188 | 135 |
| Virulence, detoxification and adaptation (IX) | 91 | 85 |
| Conserved hypothetical function (X) | 911 | 467 |
| Proteins of unknown function (XI) | 607 | 24 |
|  | 3906 | 1982 |

As Table 1 shows, there were 467 conserved proteins with hypothetical functions and 24 with unknown functions. In the next step, the functional annotation of these proteins was done using Pfam, SMART, Interpro Scan and BLASTp to predict their functional roles in *Mycobacterium tuberculosis*. Out of the 467 conserved hypothetical proteins, 259 proteins were annotated (Supplementary Table S3). No reference was found for the 24 proteins with unknown functions. Figure 3 shows the distribution of the 259 conserved hypothetical proteins into different functional classes. A large number of proteins (n) were found to be involved in intermediary metabolism and respiration (n=125 or 48%), followed by cell wall processes (n=43 or 17%), regulatory proteins (n=38 or 15%) and information pathways (n=33 or 13%). The high percentage of proteins involved in intermediary metabolism and respiration suggests that these processes are crucial for the bacteria’s energy production and survival, which could indicate its ability to sustain growth and establish infection within the host. This could also be related to its capacity for colonisation and persistence. The percentages of proteins belonging to lipid metabolism (n=12), IS elements and bacteriophages (n=4), and virulence, detoxification and adaptation (n=3), were very low. Only one protein was found from the PE and PPE protein family. No protein was found in stable RNAs.

**Figure 3:**
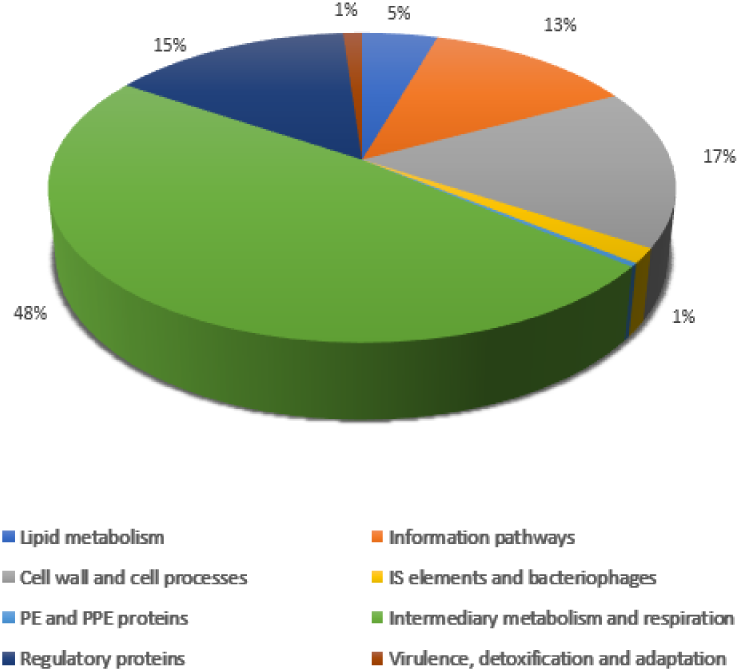
Distribution of the 259 annotated conserved hypothetical proteins into functional categories suggested by Smith (2003)

### 3.2 Reverse Vaccinology identifies 24 antigenic proteins from the 1982 conserved proteins

Figure 4 shows in summary form the outcomes of the complete reductionist reverse vaccinology process performed on the 1982 conserved proteins to reduce the time and resources needed to identify potential antigenic TB proteins as vaccine candidates.

**Figure 4:**
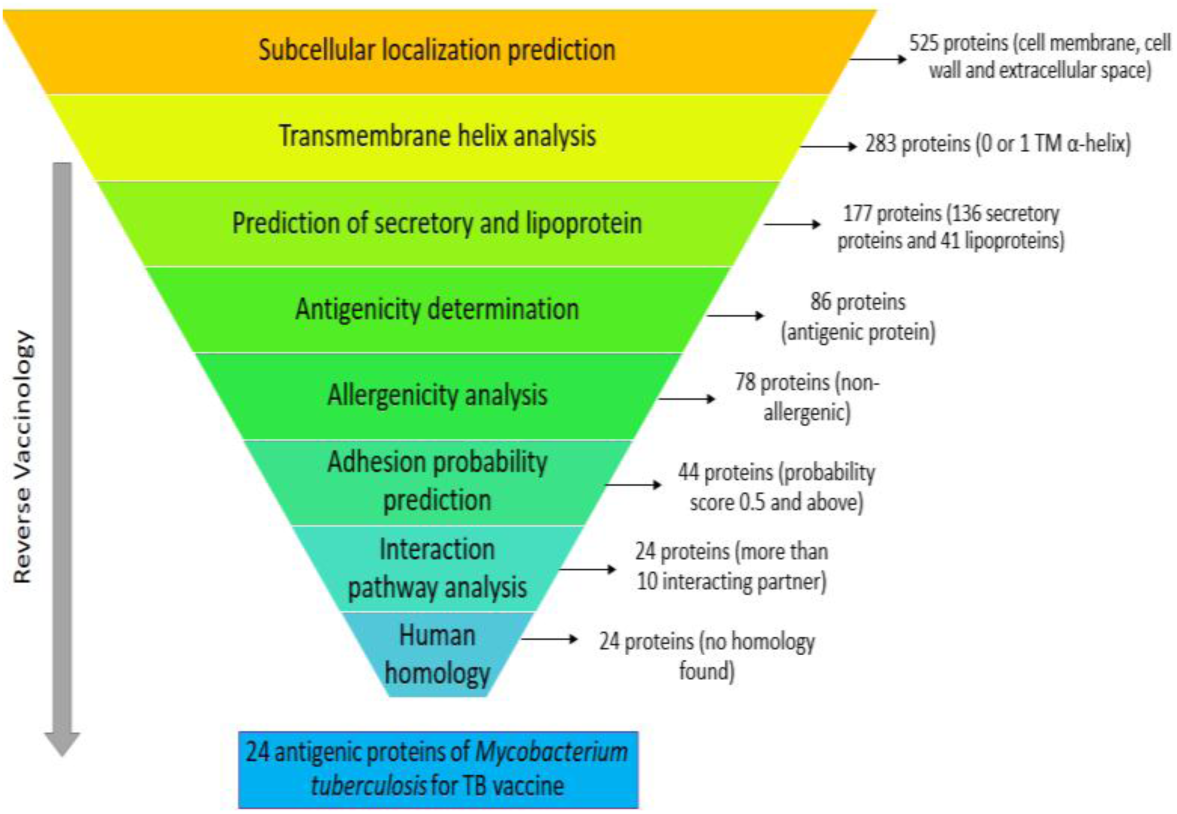
Reverse vaccinology approach used for the identification of 24 *Mycobacterium tuberculosis* antigens from 1982 conserved proteins

Subcellular localisation prediction was first undertaken to identify the location of each protein in *Mycobacterium tuberculosis.* The outer membrane and extracellular space proteins are involved in membrane integrity and permeability, efflux mechanism and active transport of molecules. They are considered suitable TB vaccine candidates since they are easily accessible to the host immune system. As predicted by four or more tools, the proteins in the cell membrane, cell wall and extracellular space of *Mycobacterium tuberculosis* were selected in this step. With the help of PSORTb v.3.0, CELLO, LocTree3, SOSUI, pLoc_bal-mGpos and GramLocEN, 525 proteins were filtered from the 1982 conserved proteins. These 525 proteins were localized in the cell membrane, cell wall and extracellular space of *Mycobacterium tuberculosis*. Next, with the help of the TMHMM server, out of the 525 proteins, 283 proteins with 0 or 1 transmembrane α-helix were found. To aid in the design of a potential vaccine, proteins with 0 or 1 transmembrane α-helix were selected. This is because proteins with multiple transmembrane α-helices present challenges in laboratory cloning and purification.

Secretory signal peptides are ubiquitous protein-sorting signals that help translocate a protein across the cytoplasmic membrane in mycobacteria and are considered essential antigenic and immunogenic TB vaccine targets. Studies have shown that *Mycobacterium tuberculosis* secretory proteins are involved in mycobacteria’s virulence [12]. The Sec pathway helps in the translocation of proteins in their unfolded state. The TAT pathway translocates proteins in their folded state. One hundred nineteen proteins were involved in the Sec pathway predicted using SecretomeP and the SignalP 4.1 server. PRED-TAT identified 17 proteins involved in the TAT pathway. Thus, of the 283 proteins, 136 were discovered to be engaged in the secretory pathway of *Mycobacterium tuberculosis*. Next, with the help of PRED-LIPO, 41 lipoproteins were identified from the 283 proteins. Lipoproteins are an essential set of membrane proteins that perform different functions in *Mycobacterium tuberculosis*, such as modulating the host’s immune response, translocation of virulent factors in the macrophages, signal transduction, and the uptake of nutrients. Thus, 177 proteins (136 secretory proteins and 41 lipoproteins) were selected for further analysis (Figure 4).

Three bioinformatics tools were used to predict antigenicity of the 177 conserved proteins. Concordance analysis by VaxiJen, VirulentPred and MP3 showed that 86 proteins were potentially highly antigenic compared to the remaining proteins (91 proteins). Algpred was used to predict the allergenic or non-allergenic proteins. From the 86 antigenic proteins, 78 proteins were found to be non-allergenic. These 78 conserved proteins were further analysed for adhesion probability using SPAAN, which identifies adhesin proteins. Adhesin proteins help attach *Mycobacterium tuberculosis* to the host’s cell-surface receptors. Forty-four proteins had a strong adhesion probability with a score of 0.5 and above. Hence, using the STRING database, these 44 proteins were chosen to study Mycobacterium tuberculosis pathway interactions. Twenty-four out of the 44 conserved and antigenic proteins were found to have more than ten interacting partners with a confidence score of 0.4. Targeting a protein with many interacting partners in the membrane region of *Mycobacterium tuberculosis* can significantly weaken the cell membrane and cell wall, thus leading to a quicker death of the bacterial cell. Following this, the prediction of the homology between humans and *Mycobacterium tuberculosis* was performed. The homologous proteins may initiate autoimmune reactions causing severe health problems in the host. No similarity was found between the 24 *Mycobacterium tuberculosis* proteins and humans.

A total of 24 conserved, membrane-spanning, antigenic, non-allergic proteins with high adhesion probability and non-homologous to humans were thus selected using the reverse vaccinology approach as shown in Table 2. An investigation into their function revealed that, out of the 24 proteins, nine proteins were observed in the cell walls and the cell processes functional class (IIII). Six proteins belonged to the PE and PPE protein families (VI). Three proteins were involved in lipid metabolism (I), whereas only one belonged to the virulence, detoxification, and adaptation class (IX).

**Table 2:**
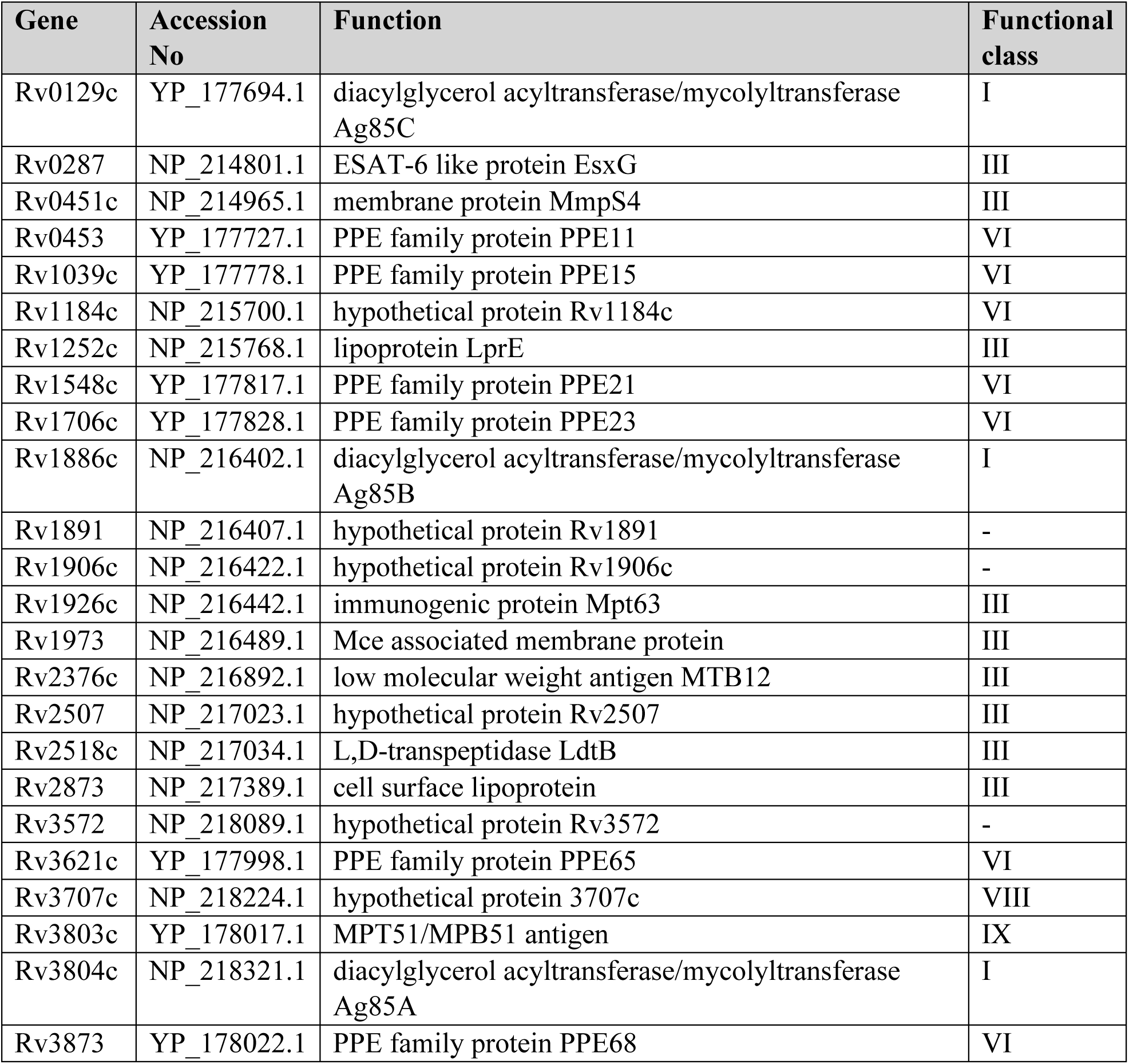
List of 24 surface-exposed, antigenic and non-allergic proteins identified by reverse vaccinology approach. Column 1-gene id, column 2-accession number of the protein, column 3-function of protein, column 4-functional class given by Smith (2003)

| Gene | Accession No | Function | Functional class |
| --- | --- | --- | --- |
| Rv0129c | YP_177694.1 | diacylglycerol acyltransferase/mycolyltransferase Ag85C | I |
| Rv0287 | NP_214801.1 | ESAT-6 like protein EsxG | III |
| Rv0451c | NP_214965.1 | membrane protein MmpS4 | III |
| Rv0453 | YP_177727.1 | PPE family protein PPE11 | VI |
| Rv1039c | YP_177778.1 | PPE family protein PPE15 | VI |
| Rv1184c | NP_215700.1 | hypothetical protein Rv1184c | VI |
| Rv1252c | NP_215768.1 | lipoprotein LprE | III |
| Rv1548c | YP_177817.1 | PPE family protein PPE21 | VI |
| Rv1706c | YP_177828.1 | PPE family protein PPE23 | VI |
| Rv1886c | NP_216402.1 | diacylglycerol acyltransferase/mycolyltransferase Ag85B | I |
| Rv1891 | NP_216407.1 | hypothetical protein Rv1891 | - |
| Rv1906c | NP_216422.1 | hypothetical protein Rv1906c | - |
| Rv1926c | NP_216442.1 | immunogenic protein Mpt63 | III |
| Rv1973 | NP_216489.1 | Mce associated membrane protein | III |
| Rv2376c | NP_216892.1 | low molecular weight antigen MTB12 | III |
| Rv2507 | NP_217023.1 | hypothetical protein Rv2507 | III |
| Rv2518c | NP_217034.1 | L,D-transpeptidase LdtB | III |
| Rv2873 | NP_217389.1 | cell surface lipoprotein | III |
| Rv3572 | NP_218089.1 | hypothetical protein Rv3572 | - |
| Rv3621c | YP_177998.1 | PPE family protein PPE65 | VI |
| Rv3707c | NP_218224.1 | hypothetical protein 3707c | VIII |
| Rv3803c | YP_178017.1 | MPT51/MPB51 antigen | IX |
| Rv3804c | NP_218321.1 | diacylglycerol acyltransferase/mycolyltransferase Ag85A | I |
| Rv3873 | YP_178022.1 | PPE family protein PPE68 | VI |

The literature has shown that these proteins (Table 2) are essential in escaping the human immune response. PPE11 is involved in the pathogenesis of TB and macrophage persistence [76] and PPE15 inhibits the production of reactive nitrogen inside alveolar macrophages [77]. Mazandu and Mulder (2012) predicted the role of PPE21 in the pathogenicity of *Mycobacterium tuberculosis* [78]. PPE65 helps evade the adaptive immune response by obstructing the helper-T cell response [79]. ESAT-6, belonging to cell wall and cell processes functional class, regulates macrophage apoptosis [80]. The membrane protein, MmpS4, plays an essential role in the growth of *Mycobacterium tuberculosis* under iron-deficiency conditions [81]. Goulding et al. (2009) suggest that immunogenic protein Mpt63 is involved in the virulence mechanism inside the host body [82]. A recombinant vaccine constructed using Ag85A and Ag85B involved in lipid metabolism is in phase-I of a TB vaccine clinical trial [1]. Thus, the literature validates the selected antigenic proteins as some of the most crucial proteins required by the *Mycobacterium tuberculosis* survival mechanism. Therefore, constructing an epitope-based TB vaccine from these 24 proteins would help generate a robust immune response in humans.

### 3.3 Immunoinformatic analysis for the prediction of B-cell and T-cell epitopes for TB vaccine construction

Epitopes are the part of an antigen that binds to T-cells, B-cells, or antibodies in the host body. The T-cells and B-cells are the principal constituents of the immune system. For predicting epitopes, immunoinformatics analysis is performed to predict the B-cell and T-cell (MHC-I and II-restricted) epitopes and then the most suitable vaccine candidates are filtered based on immunogenicity, toxicity allergenicity and hydrophilicity.

#### 3.3.1 B-cell epitopes for TB vaccine

B-cells are essential in providing humoral or antibody-mediated responses by producing antibodies that can neutralize *Mycobacterium tuberculosis* or facilitate its clearance by other immune cells through processes such as opsonization. Epitopes recognised by the antibody’s paratopes (antigen/epitope-recognition site in antibodies) or B-cell receptors (BCR) are termed B-cell epitopes. Bepipred, ABCpred and BCPREDS were used identify the B-cell epitopes. From all three bioinformatics tools, 280 B-cell epitopes were identified from the 24 antigenic proteins. These 280 epitopes had varying lengths of 2-40 amino acid residues. A B-cell epitope with 8-20 residue size was considered a suitable vaccine candidate because this range is optimal for ensuring adequate interaction with B-cell receptors (BCRs), enabling effective antibody production without compromising epitope stability. Concordance analysis was undertaken to identify B-cell epitopes commonly predicted by all three tools to improve prediction accuracy. 74 B-cell epitopes of 8-20 residues in length were common among these 24 antigenic TB proteins.

After selecting the 74 B-cell epitopes, filtering was performed to investigate safe and immunodominant B-cell epitopes for vaccine construction (Supplementary Table S4). First, the immunogenic nature of B-cell epitopes was predicted using the VaxiJen server and ToxinPred. 38 non-toxic B-cell epitopes with a Vaxijen score higher than 0.8 were selected for further analysis. The selection of hydrophilic epitopes for constructing a TB vaccine helps in swift interactions between a vaccine and the host cells. This enables a high chance of initiating a robust immune response in the host. The hydrophilicity prediction was made using the ProtParam server to calculate the GRAVY score. This study excluded B-cell epitopes with positive GRAVY scores, indicating hydrophobicity, to prioritise hydrophilic epitopes that enhance vaccine efficacy. From ProtParam analysis, 9 B-cell epitopes with negative GRAVY scores were selected. The ALLERTOP 2.0 bioinformatics tool was used to identify non-allergenic epitopes further. Out of 9 B-cell epitopes, one epitope was found to be allergenic and excluded from the study. At the end of the immunoinformatics analysis, 8 B-cell epitopes were shortlisted for TB vaccine construction.

#### 3.3.2 MHC-I restricted T-cell epitopes for TB vaccine

T-cells play a significant role in cell-mediated or cellular immunity. In cellular immunity, *Mycobacterium tuberculosis* is ingested by antigen-presenting cells, such as macrophages, dendritic cells, etc. The pathogen is then fragmented into smaller antigenic peptides. These are then presented to T-cell receptors (TCR) on the surface of T-cells through the cell-surface attached major histocompatibility complex (MHC) molecule. Most T-cell epitope prediction tools are trained with experimentally validated 9-mer epitope residues and 15-mer epitope residues for MHC-I and MHC-II restricted epitope prediction, respectively. The epitopes binding to MHC-I molecules are cytotoxic T-cell (CTL) epitopes or MHC-I restricted T-cell epitopes. The CTL epitope was set at nine amino acid residues. CTL epitopes were predicted using IEDB MHC-I, NetCTLpan 1.1 and IEDB MHC-NP. From the IEDB server analysis, CTL epitopes with higher binding affinity and a percentile rank lower than 0.4 were selected. The 24 TB antigens had 469 CTL epitopes from all three servers. Two hundred and seven CTL epitopes were found to be commonly predicted by all three servers.

To confirm the ability of CTL epitopes to initiate an immune response, antigenicity and toxicity analysis were performed using VaxiJen and Toxin Pred. Supplementary Table S5 shows the results of antigenicity, toxicity, hydrophilicity, and allergenicity. The antigenicity and toxicity scores revealed that out of 207 CTL epitopes, 58 epitopes were highly immunogenic and non-toxic. These 58 epitopes were considered for further hydrophilic score calculations and, out of these, 25 CTL epitopes were discovered to be hydrophilic, as indicated by their negative GRAVY scores. The allergenic nature of the epitopes was then determined using ALLERTOP 2.0. Out of 25 CTL epitopes, 11 were found to cause allergenic reactions inside the host body. These epitopes were excluded from the study. Finally, 14 CTL epitopes were chosen for constructing an epitope-based TB vaccine (Supplementary Table S6).

#### 3.3.3 MHC-II restricted T-cell epitopes for a TB vaccine

MHC-II molecules are present on macrophage and dendritic cells. The primary function of MHC-II is to present antigens/epitopes to the naïve T-helper cells. This interaction leads to the release of cytokines that help the development of naïve T-helper cells into effector or memory T-cells. The epitopes presented by MHC-II molecules are termed helper T-cells (HTL epitopes) or MHC-II restricted T-cell epitopes. The length of HTL epitopes was set at 15 amino acid residues. IEDB MHC-II and the NetMHCIIpan 3.2 server identified 428 epitopes in the 24 antigen proteins of *Mycobacterium tuberculosis*. Concordance analysis estimated 346 common HTL epitopes between the two methods. No common epitope was found between the two methods for one protein with accession number NP_214965.1. Out of the selected 346 HTL epitopes, 250 were found to be immunogenic and non-toxic, after rigorous antigenicity and toxicity analysis using VaxiJen and ToxinPred. Supplementary Table S7 shows the antigenicity, toxicity, hydrophilicity and allergenicity analysis outcomes. Only six of the 250 immunogenic and non-toxic HTL epitopes were hydrophilic. All remaining HTL epitopes were excluded from the study. Finally, five HTL epitopes remained after evaluating the allergenic nature of epitopes.

In summary, after selecting highly immunogenic and excluding cross-reactive epitopes, 27 epitopes (8 B-cell epitopes, 14 CTL epitopes and 5 HTL epitopes) were found from 18 (out of 24) antigenic *Mycobacterium tuberculosis* proteins (Supplementary Table S8).

The selected 27 epitopes from the 18 proteins that can evoke a potent cellular and humoral immune response in the host are listed in Table 3. Three proteins were found to have both B-cell and T-cell epitopes. No suitable vaccine candidates were found in six proteins: NP_214801.1, YP_177828.1, NP_217034.1, YP_178017.1, YP_178022.1 and NP_218089.1. Out of these six proteins, the functional significance of three proteins (NP_217034.1, NP_218089.1 and YP_178017.1) was unknown in *Mycobacterium tuberculosis*. The selected epitopes are expected to evoke a strong and specific immune response to provide broad immune protection against a large number of *Mycobacterium tuberculosis* strains.

**Table 3:**
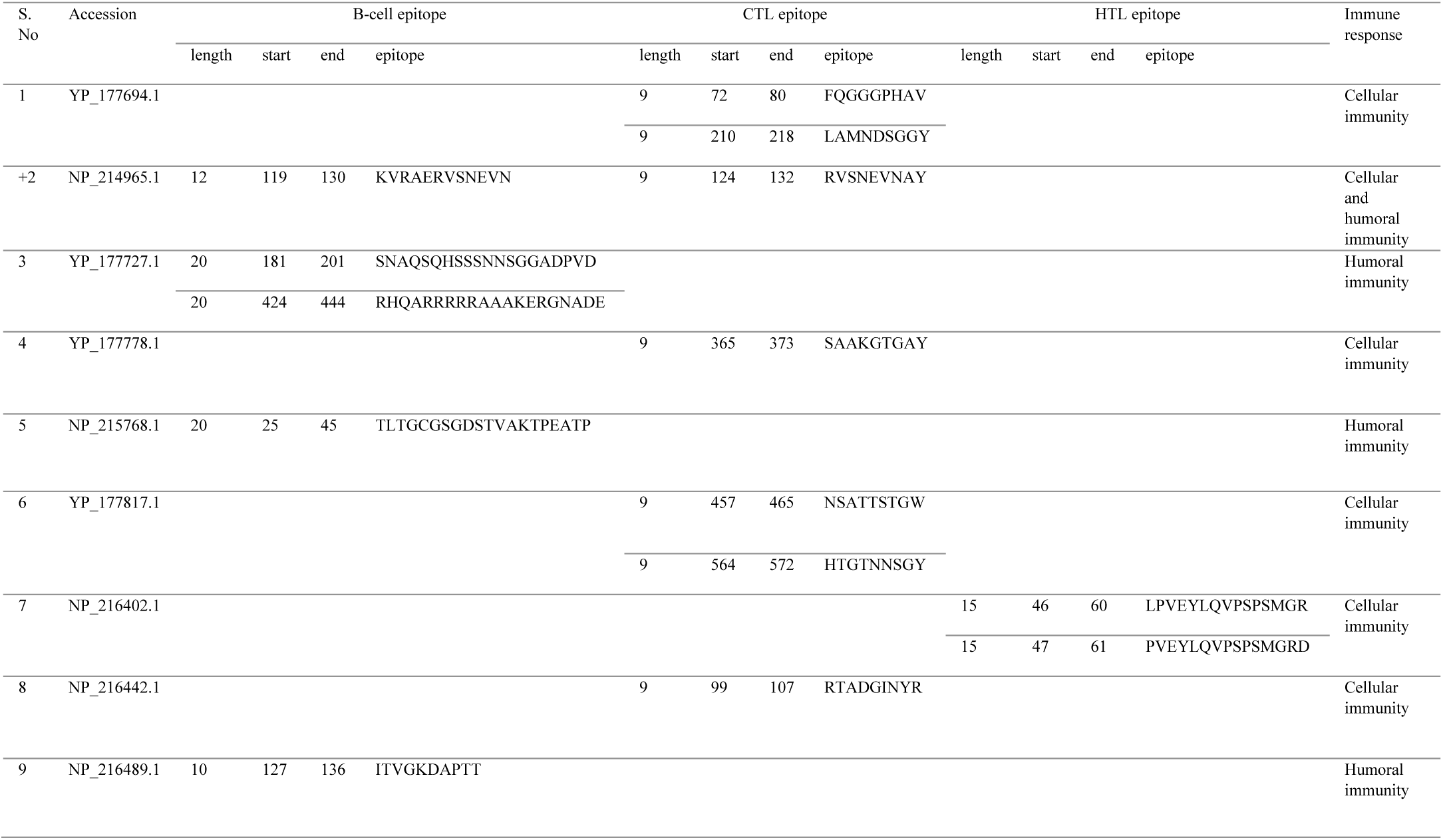
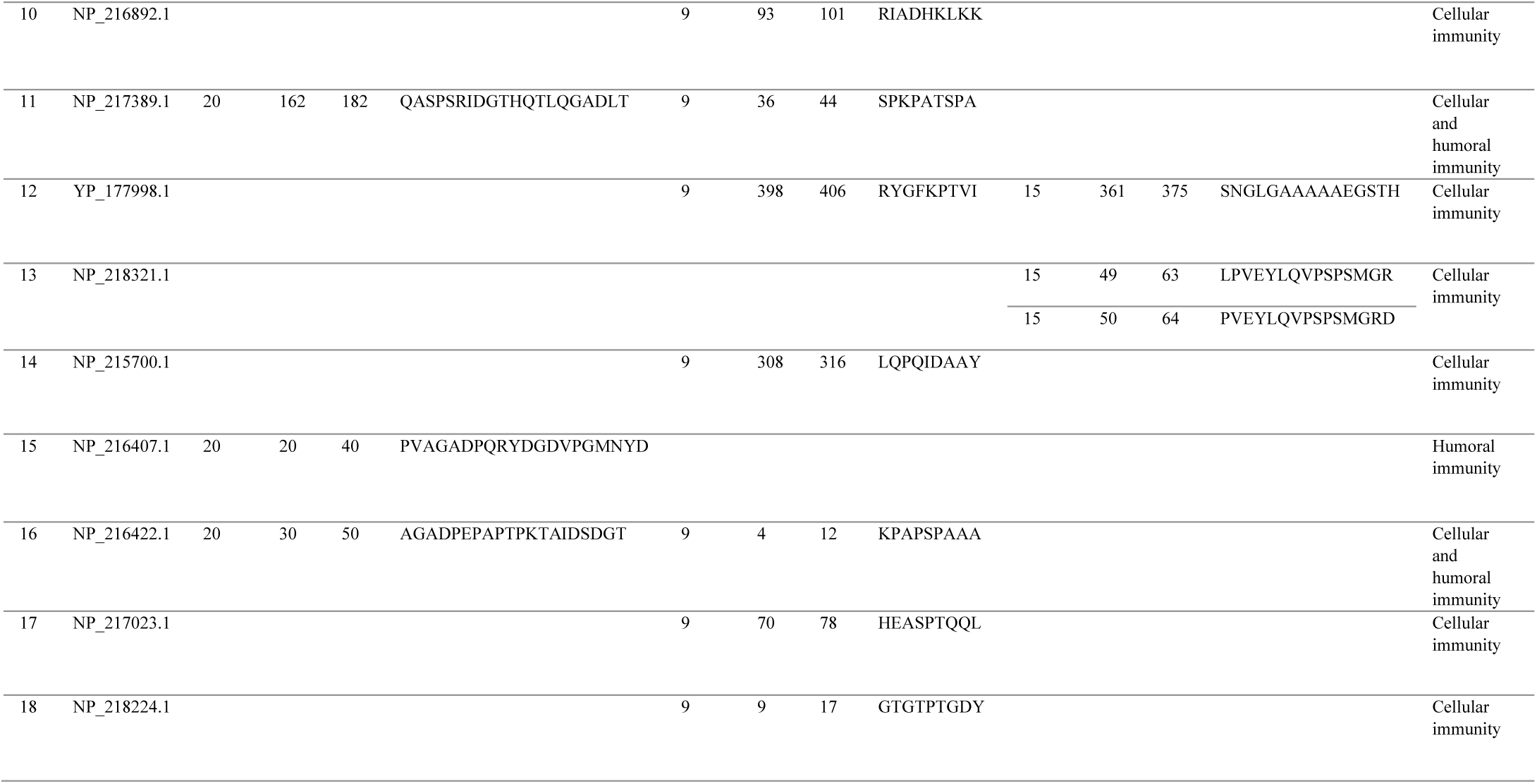
Final 27 epitopes (8 B-cell epitopes, 14 CTL epitopes and 5 HTL epitopes) selected from 18 *Mycobacterium tuberculosis* antigens for constructing an epitope-based vaccine for Tb. Length represents the length of an epitope or number of amino acid residues in an epitope, start and end represents starting and ending position of an epitope in the protein, and epitope represents letter code for amino acid residues.

#### 3.3.4 Population coverage analysis

MHC molecules are highly polymorphic, and different HLA types are expressed at different rates in different ethnicities around the world. Therefore, to design a universal vaccine against TB, there is a need to analyse population coverage of the selected T-cell epitopes to minimize the risk of developing an ethnically biased vaccine [57]. The prevalence of MHC-I and II HLA alleles in diverse ethnic groups of the world help determine the most prevalent set of T-cell epitopes in global population. Thus, the population coverage assessment of the predicted CTL and HTL epitopes and their corresponding MHC-I and MHC-II HLA alleles was done using IEDB epitope analysis resource (IEDB-AR). Among 15 regions, the results exhibited maximum population coverage in Europe (99.74%), closely followed by North America (99.54%), East Asia (99.44%), West Indies (98.14%), South Asia (97.09%), Oceania (96.94%), Southeast Asia (96.52%), Northeast Asia (95.53%), North Africa (95.06%) and others (Figure 5). The lowest coverage was found in South Africa (83.76%) that followed Central Africa (89.09%). As a result, 99.16% of the world’s population was covered by the predicted CTL and HTL epitopes from the 18 antigens of *Mycobacterium tuberculosis*. This analysis reveals that the strategy of developing a universal TB vaccine from the selected epitopes could be successful and highly effective in global eradication or suppression of TB.

**Figure 5:**
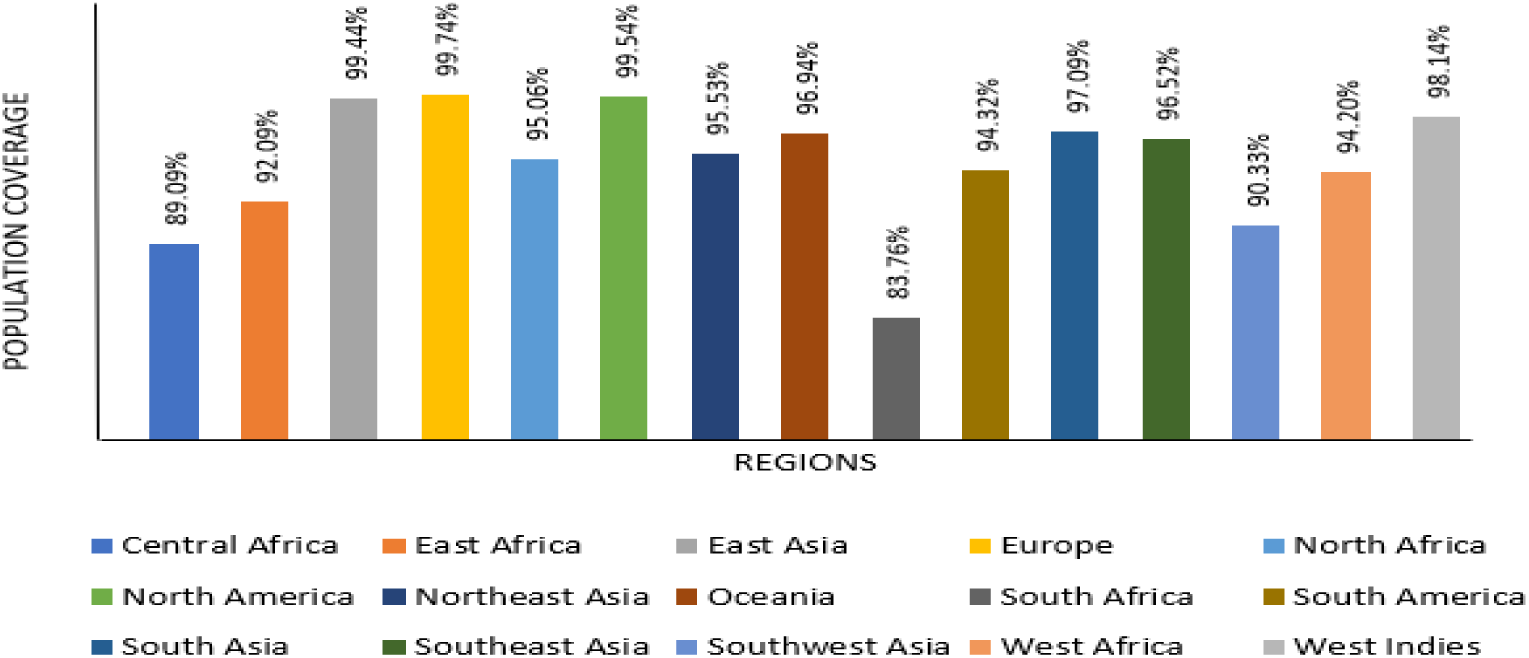
Population coverage analysis predicted by IEDB-AR for 14 CTL epitopes and 5 HTL epitopes with their respective MHC HLA alleles. The percentage of population coverage was calculated for 15 regions covering the globe

### 3.4 Designing a structural model of the proposed TB vaccine

After selecting the final TB vaccine epitopes, the vaccine sequence was designed, and the vaccine structural model was constructed to determine its efficacy of interaction inside the host.

#### 3.4.1 Vaccine sequence design

The selected 27 epitopes shown in Table 4 were used for designing the TB vaccine protein sequence.

**Table 4:** Final 27 epitopes for TB vaccine construction.

| Epitope No. | CTL epitopes (E) | Epitope No. | HTL epitopes(E) | Epitope No. | B-cell epitopes(E) |
| --- | --- | --- | --- | --- | --- |
| 1 | FQGGGPHAV | 15 | LPVEYLQVPSPSMGR | 20 | KVRAERVSNEVN |
| 2 | LAMNDSGGY | 16 | PVEYLQVPSPSMGRD | 21 | SNAQSQHSSSNNSGGADPVD |
| 3 | RVSNEVNAY | 17 | SNGLGAAAAAEGSTH | 22 | RHQARRRRRAAAKERGNADE |
| 4 | SAAKGTGAY | 18 | LPVEYLQVPSPSMGR | 23 | TLTGCGSGDSTVAKTPEATP |
| 5 | NSATTSTGW | 19 | PVEYLQVPSPSMGRD | 24 | ITVGKDAPTT |
| 6 | HTGTNNSGY |  |  | 25 | QASPSRIDGTHQTLQGADLT |
| 7 | RTADGINYR |  |  | 26 | PVAGADPQRYDGDVPGMNYD |
| 8 | RIADHKLKK |  |  | 27 | AGADPEPAPTPKTAIDSDGT |
| 9 | SPKPATSPA |  |  |  |  |
| 10 | RYGFKPTVI |  |  |  |  |
| 11 | LQPQIDAAY |  |  |  |  |
| 12 | KPAPSPAAA |  |  |  |  |
| 13 | HEASPTQQL |  |  |  |  |
| 14 | GTGTPTGDY |  |  |  |  |

The docking analysis of the 27 filtered epitopes with every other epitope created 153 combinations of various epitopes. The best combination of 27 epitopes with a strong binding affinity to one another, was used for constructing vaccines. The structure of the PADRE sequence and the 27 epitopes attached with the linker was constructed using PEPstrMOD. First, the affinity of the PADRE sequence with the five HTL epitopes was determined using PatchDock and FireDock. The PADRE-E17 (HTL epitope) was the best combination compared to the other four HTL epitopes based on lowest binding energy (Supplementary Table S9). The structure of PADRE-E17 was then constructed using the I-TASSER server. The binding compatibility of PADRE-E17 was determined for the remaining HTL epitopes. E-15 was found to have strong binding to PADRE-E17. The structure of the PADRE-E17-E15 combination was then generated using I-TASSER. This process was first undertaken for the HTL epitopes, followed by CTL epitopes and then the B-cell epitopes. Figure 6 (i) shows the final TB vaccine sequence obtained through the extensive analysis of the TB epitope combinations. The adjuvant 50S ribosomal protein L7/L12 at the N-terminal and β-defensin at the C-terminal were added to the vaccine protein sequence with the help of the EAAAK linker. With the two adjuvants and linkers, the final length of the TB vaccine sequence was 629 amino acid residues (Figure 6 (ii)).

**Figure 6:**
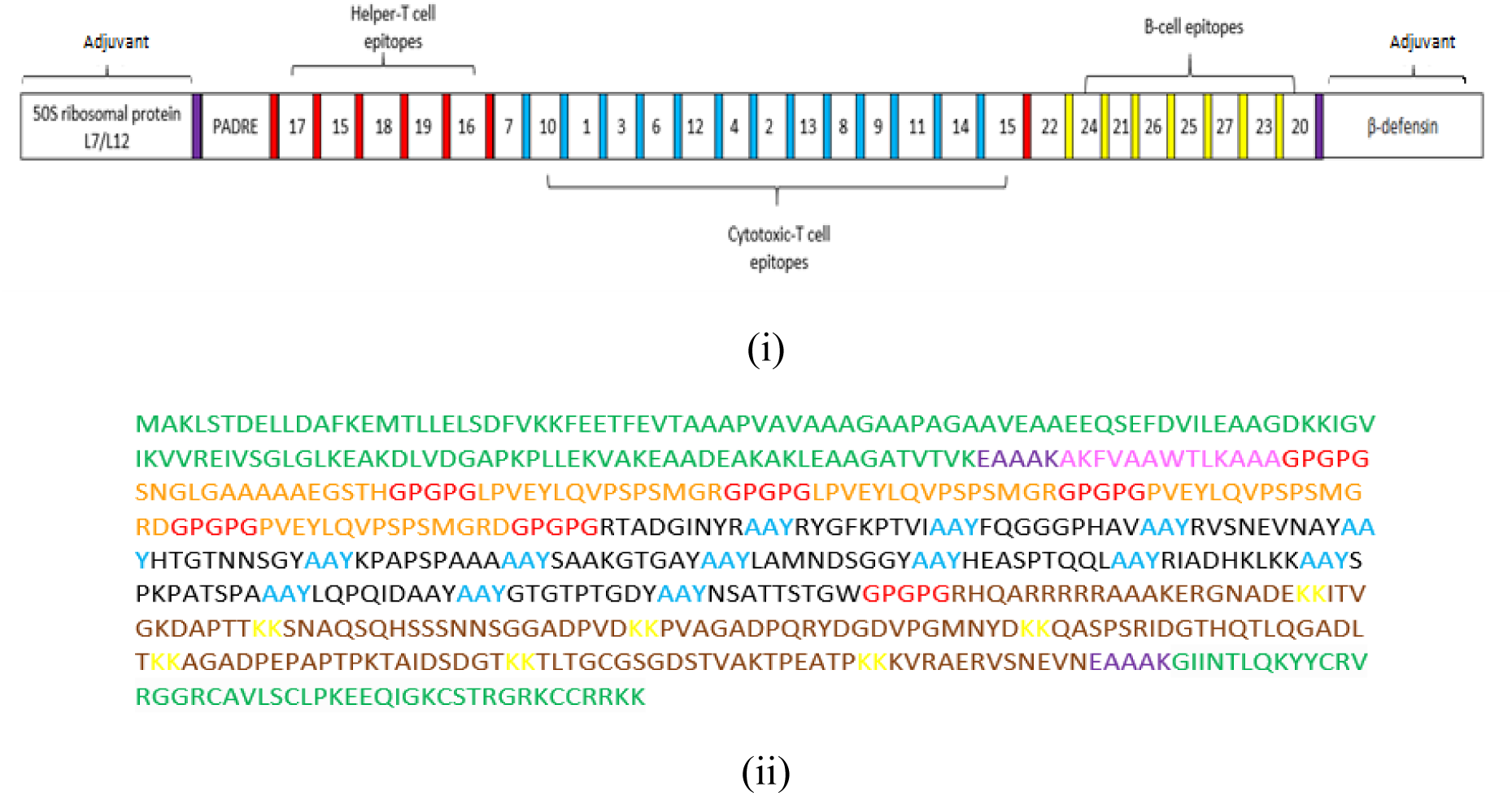
Epitope-based vaccine sequence construction scheme: (i) Schematic representation of TB vaccine construct consisting of 27 B-cell and T-cells epitopes joined by flexible linkers and adjuvants at N- and C-terminals of the vaccine protein. The epitope combinations were found after determining their binding affinity using PatchDock and FireDock, and (ii) TB vaccine protein sequence. The adjuvant sequence is highlighted in green, the PADRE sequence in pink, and the flexible linkers in purple (EAAAK), red (GPGPG), blue (AAY) and yellow (KK). The HTL epitopes are highlighted in orange, the CTL epitopes are black, and the B-cell epitopes are brown

#### 3.4.2 Assessment of the properties of the TB vaccine construct

After designing the TB vaccine sequence, its antigenicity, allergenicity and physiochemical properties were evaluated. The antigenicity was predicted using the VaxiJen server. The high antigenic score of 0.9126 showed the great immunogenic potential of the constructed TB vaccine. The AlgPred server indicated that the vaccine designed is non-allergenic to the host. ProtParam was used for predicting the physiochemical properties of the designed TB vaccine sequence. Proteins with MW less than 100 kDa indicate the possibility of experimentally studying the proteins in the laboratory for vaccine or drug development. Thus, proteins with a molecular weight of more than 100 kDa were not considered potential vaccine targets.

The vaccine sequence comprised 629 amino acid residues with a molecular weight of 64.86 kDa. The TB vaccine construct contained 9071 atoms, and its molecular formula was C_2837_H_4507_N_817_O_895_S_15._ The isoelectric point (pI or pH(I)) is the pH at which a molecule carries no net electrical charge. The pI was assessed to be 9.19, indicating the vaccine construct was slightly basic. This characteristic is significant as it suggests that the vaccine construct will have a net positive charge under physiological conditions, which could enhance its interaction with negatively charged cell membranes and potentially improve its stability and effectiveness in eliciting an immune response. *In vivo* half-life predicts the time for half the amount of protein to disappear within a cell. A half-life between 2 to 100 hours is considered suitable for protein degradation, depending on the nature of the protein. The estimated half-life of the TB vaccine protein was 30 hours in mammalian reticulocytes (*in-vitro*). An instability index with a value of 37.94 represented the stable nature of the TB vaccine. The high value of the aliphatic index (62.29) indicated the high thermostability of the protein. The GRAVY score of the vaccine was found to be -0.503 where the negative value indicated the hydrophilic nature of the vaccine. The designed TB vaccine thus had all the characteristics of a promising vaccine candidate required for initiating an effective immune response inside the host and with a global coverage.

#### 3.4.3 Structural model of the TB vaccine

Secondary structure is crucial for maintaining the three-dimensional structural stability of a protein. The secondary structure of the TB vaccine was constructed using PSIPRED (Figure 7). According to PSIPRED, 213 amino acid residues formed alpha-helixes, constituting 33.86% of the overall TB vaccine sequence. Only 66 residues participated in beta-strand formation (10.49%), whereas 350 amino acid residues formed the coils (55.64%) in the TB vaccine sequence. High flexibility in a protein’s secondary structure is indicated by the proportion of its structural elements—alpha helices, beta strands, and coils. In the context of the TB vaccine construct, high flexibility is associated with a higher proportion of coil regions (55.64%) compared to alpha helices (33.86%) and beta (10.49%) strands suggesting high flexibility of the TB vaccine protein, facilitating smoother interactions with immune cells.

**Figure 7:**
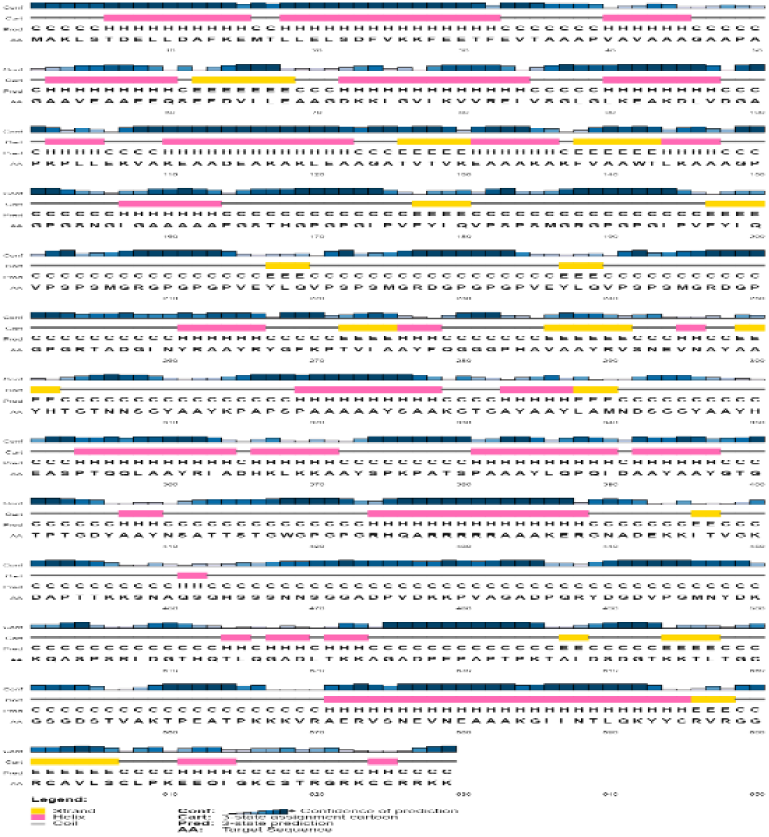
Secondary structure prediction of the TB vaccine sequence using PSIPRED. The sequence comprised alpha-helixes (33.86%) (pink), beta-strands (10.49%) (yellow) and coils (55.64%) (light grey) indicating that the constructed vaccine is highly flexible

The tertiary structural model of the TB vaccine was constructed using the RaptorX server. RaptorX performed homology modelling, and 1DD3_A PDB entry was used as a template for generating a structural model. All 629 amino acid residues were modelled, with only 10% being in the disordered region. In the constructed model, 61% of the residues were exposed, 17% were medium, and 21% were buried in folded conformation. Before refining the structural model, the Ramachandran plot showed that 87.6% of the amino acid residues were in the most favoured region. Thus, the structural model was refined using the GalaxyRefine server to improve the quality of the predicted structural model. Figure 8 (i) shows the 3D structural model of the TB vaccine after performing the refinement step. The refined structural model had 92.3% amino acid residues in the most favoured region (Figure 8 (ii)). The parameters evaluated after refinement were GDT-HA score of 0.9112, MolProbidity of 2.027, a clash score of 10.7, an RMSD of 0.52 and poor rotamers with a value of 0.7. GDT-HA scores range from 0 to 1, with values closer to 1 indicating better backbone quality. Therefore, a score of 0.9112 implies that the backbone of the model is of high quality. MolProbity score indicates the log-weighted combination of clash score, percentage of not favoured amino acid residues in Ramachandran plot and percentage of bad sidechain rotamers [85]. MolProbity scores are lower when the model has fewer clashes and better conformational geometry. Hence, a score of 2.027 supports the conclusion that the model is of good structural quality.

**Figure 8:**
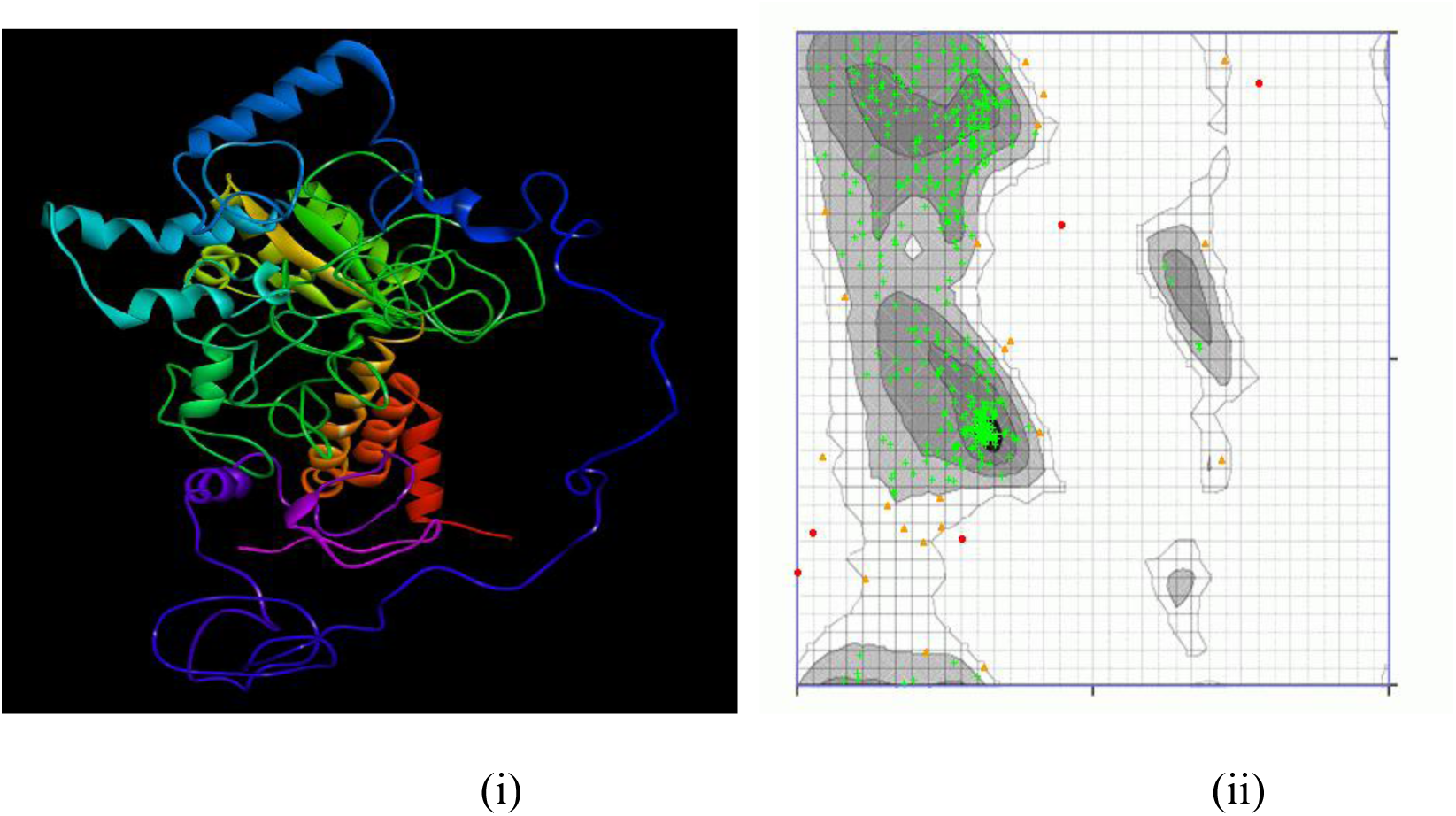
Refined structural model of the TB vaccine construct: (i) The 3D structure is coloured in rainbow colour from violet to red (from N-terminus to C-terminus), and (ii) the Ramachandran plot of the refined structure shows 92.3% amino acid residues in the most favoured region (green crosses)

#### 3.4.4 Interaction of TB vaccine construct with toll-like receptors

The epitopes present on the surface of the pathogen interact with toll-like receptors (TLRs) on the surface of the APCs, causing activation of the adaptive immune system. The strong binding of both molecules will initiate the process of immune response generation. To understand the interaction of the TB vaccine with TLRs, molecular docking was performed. The ClusPro 2.0 server performed molecular docking between the TB vaccine construct and TLR-2, TLR-4, and TLR-6 with PDB id 2Z7X, 4G8A and 4OM7, respectively. The output of the docking process displayed 30 clusters (0-29) for each docked complex. Cluster 0 of TLR-2 and the TB vaccine docked complex was found to have the lowest binding energy of -1172.8 kcal/mol. This cluster involved the highest number of members in the docking interaction, *i.e*., 66 amino acid residues of the vaccine interacting with TLR-2. Cluster 2 of the TLR-4 docked complex had the lowest binding energy of -1241.1 kcal/mol, with 52 residues involved in the interaction. For TLR-6 and the TB vaccine, cluster 0 had the lowest binding affinity of -1002.2 kcal/mol, with 38 residues involved in the docking interaction. Figure 9 shows the docked complexes and their interacting residues for the three TLRs-TB vaccine constructs. The interaction of TLR-2 and TB vaccine construct showed the presence of 21 hydrogen bonds, 7 salt bridges, and 228 non-bonded contacts. In the docked complex of TLR-4 and TB vaccine construct, 23 hydrogen bonds, 7 salt bridges and 343 non-bonded contacts were found. For TLR-6 and TB vaccine construct, 17 hydrogen bonds, 5 salt bridges, and 176 non-bonded contacts were evident. The docking results revealed that the designed TB vaccine had substantial interaction with the selected TLRs and would accomplish the goal of initiating the best immune response.

**Figure 9:**
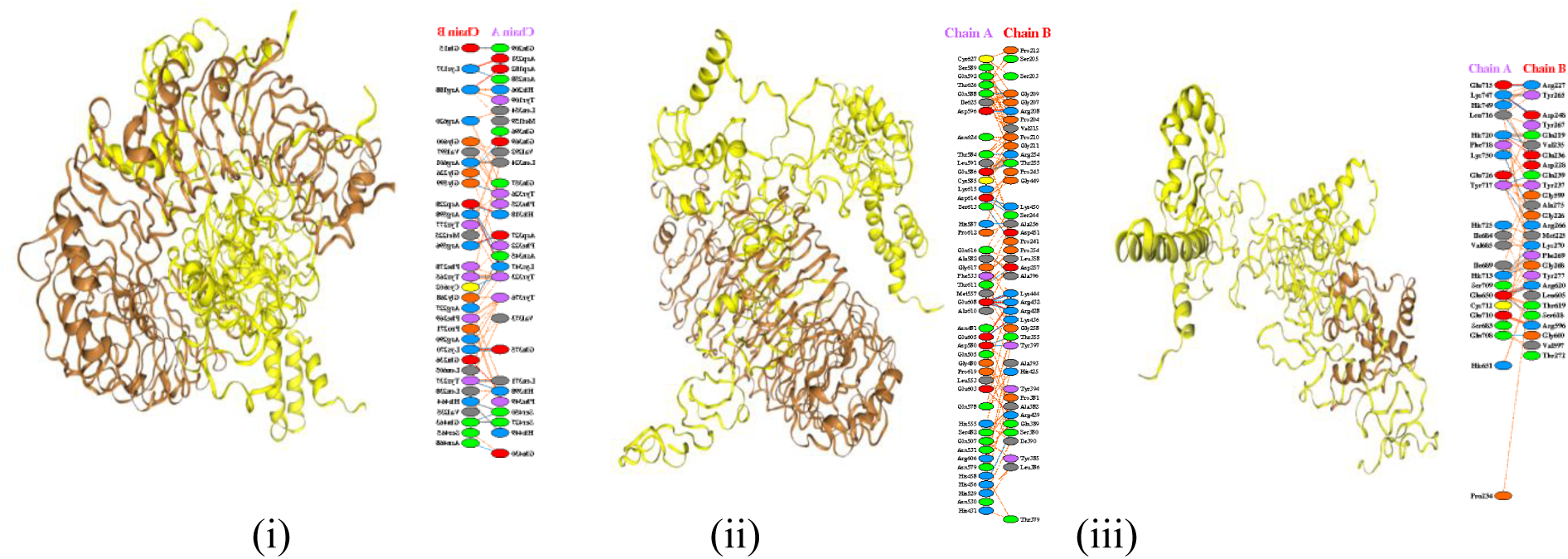
The docked complex of toll-like receptors (brown colour) with the TB vaccine construct (yellow colour). TB vaccine construct (Chain B) docked with: (i) TLR-2 (Chain A), (ii) TLR-4 (Chain A) and (iii) TLR-6 (Chain A)

Normal mode analysis (NMA) was carried out to determine the stability of the docked complex in relation to the mobility (or movement) of the vaccine residues. The iMODS server was used to perform a molecular dynamic simulation of the docked complex of the vaccine with TLR-2, TLR-4 and TLR-6. Deformation in the docked complex of the TB vaccine and TLR will delay the activation of the immune system. Deformability is the ability of amino acid residues to deform at a particular position within the protein. Analysis of the trajectory of the docked complex helps determine the occurrence of any deformation in the complex. The eigenvalue provided by the docked complex represents the energy required for deforming the complex. The eigenvalues found for the complex of TB vaccine with TLR-2, TLR-4 and TLR-6 were 1.018829e-5, 1.304754e-05 and 2.501926e-5, respectively (Figure 10). The resulting small eigenvalues indicated a low chance of deformation for the TB vaccine-TLR docked complexes. Thus, this analysis supports that our designed TB vaccine has the potential to generate a quick immune response.

**Figure 10:**
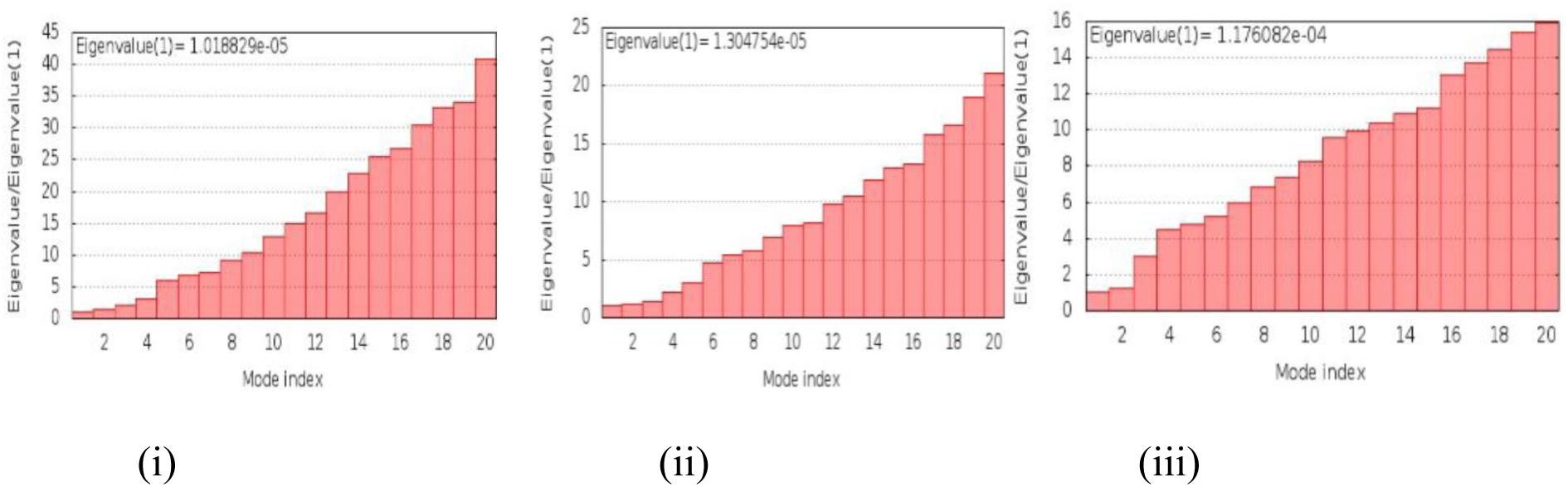
Results of molecular dynamic simulation performed using iMODS. The eigenvalue of the docked complex of TB vaccine and toll-like receptors: (i) TB vaccine-TLR2, (ii) TB vaccine-TLR4, and (iii) TB vaccine-TLR6, show low chance of deformation of the complex and high potential to generate a quick immune response.

#### 3.4.5 Codon optimisation and *in-silico* cloning of the TB vaccine

Codon optimisation is crucial because different organisms prefer specific codons to encode the same amino acid. By optimising the codon usage in the vaccine construct to match the host organism’s preference, the efficiency of protein expression is significantly improved, leading to higher yields of the vaccine protein in the chosen expression system, such as *E. coli*. JCat was used for codon optimisation of the TB vaccine construct in the *E. coli* strain K12. The length of the optimised codon sequence was observed to be 1887 nucleotides. The GC content and codon adaptation index (CAI) of the TB vaccine construct were 54.48% and 0.99, respectively. GC content and CAI signify high expression of the vaccine protein. The nucleotide sequence of the TB vaccine was then cloned using the SnapGene tool. Figure 11 shows the in-silico cloning of the TB vaccine construct. The cloning was done using a pET-28a (+) expression vector with restriction sites EcoRI and BamHI, respectively. These enzymes cut the DNA at specific recognition sequences, creating “sticky ends” that facilitate the insertion of the optimised TB vaccine construct into the vector (Figure 11). The use of two different restriction sites ensures that the insert is cloned in the correct orientation, which is critical for proper protein expression. The size of the final clone was observed to be 7256 base pairs. This size suggests that the vector can accommodate the insert without stability issues, and the cloning strategy is compatible with the vector’s replication and expression capabilities in *E. coli*. The successful in-silico cloning of the optimised TB vaccine construct into the pET-28a (+) vector indicates a high potential for successful expression in *E. coli* and enhances the likelihood that the TB vaccine protein will be produced efficiently.

**Figure 11:**
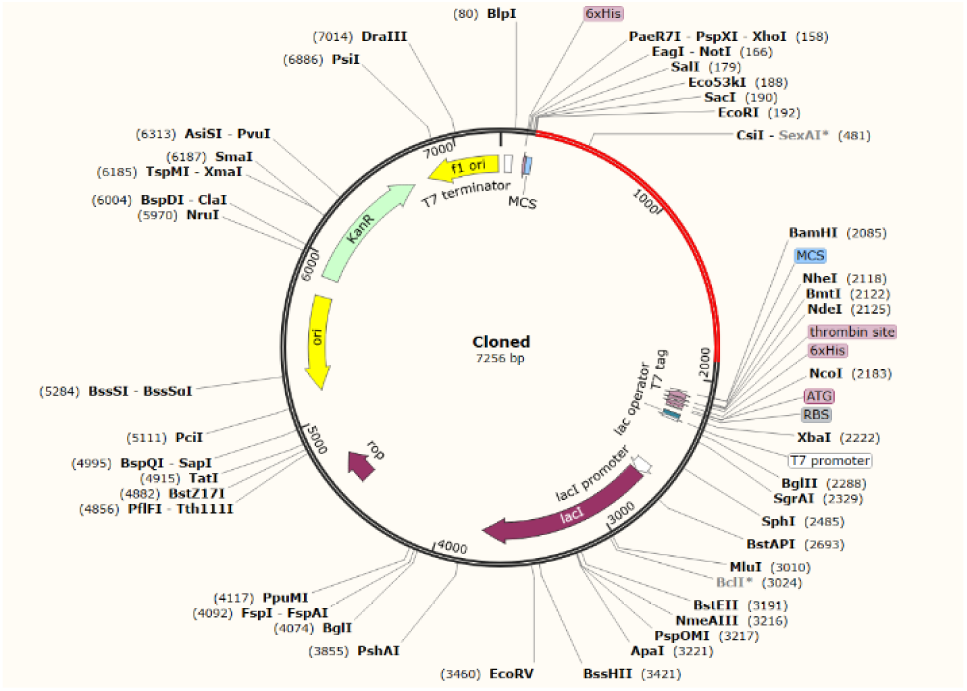
In-silico cloning of the TB vaccine using pET-28a (+) expression vector. The red part in the circle represents the codons corresponding to the designed TB vaccine, and the black represents the pET-28a (+) expression vector

### 3.5 Study of the immune response profile of the TB vaccine construct for predicting vaccine efficacy

Vaccines are cost-effective pharmaceutical products that have played an essential role in eliminating and eradicating infectious diseases. The main goal of this study was to develop a cost-effective epitope-based TB vaccine that can provide long-lasting protection and herd immunity, reduce mortality rates, fight against antimicrobial resistance and enhance the speed and strength of the host immune response against *Mycobacterium tuberculosis.* In the previous step, an epitope-based TB vaccine was designed and its beneficial properties and effective interaction with antigen-presenting cells were elucidated. It is also essential to determine the efficacy of our epitope-based TB vaccine by studying the immune response it generates. C-ImmSim server was used to examine the immune response profile of the TB vaccine construct. The vaccine construct was administered three times at four-week intervals and its impact on both humoral and cell-mediated immunity was assessed.

After exposure to the vaccine construct, the immune response generated by the host immune system was higher after every dose. Figures 12(i), 12(ii), and 12(iii) illustrate the humoral immune response following each vaccine dose.

**Figure 12:**
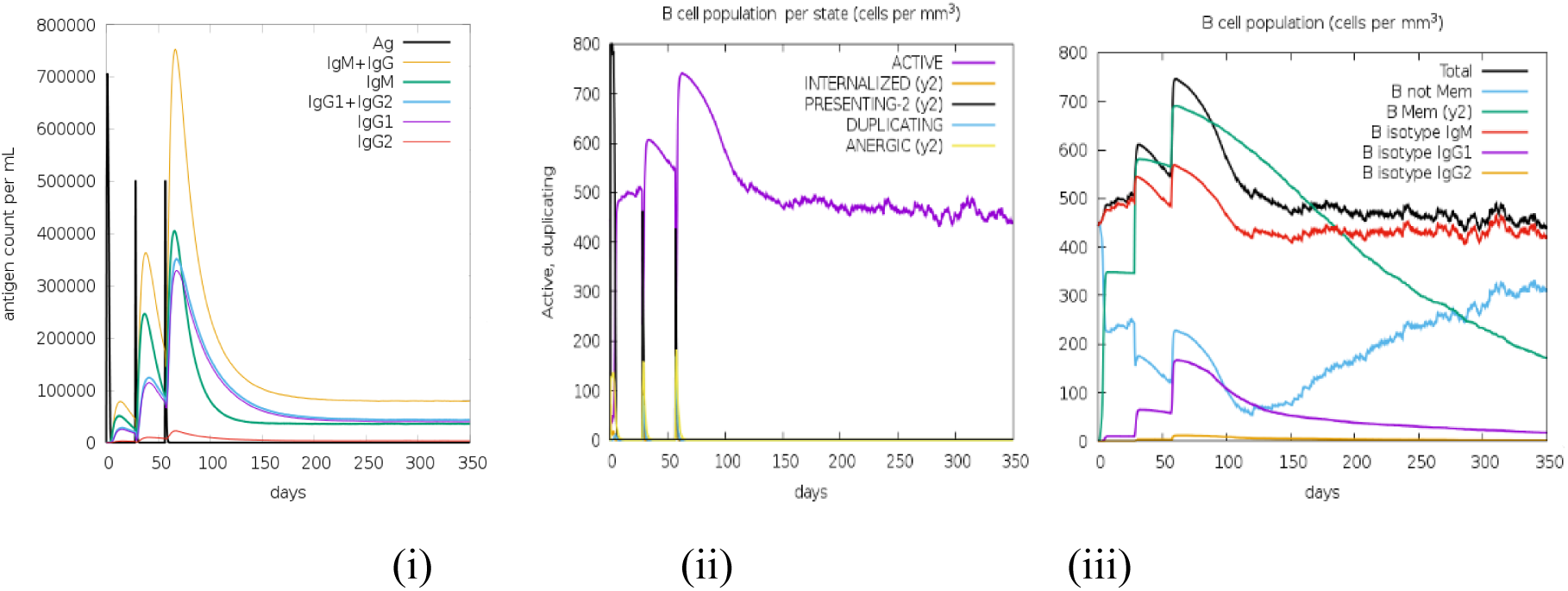
Humoral immune response profile of the epitope-based TB vaccine construct: (i) Generation of immunoglobulins upon exposure to TB vaccine construct, (ii) Number of active B-cell populations per state of antigen exposure, and (iii) Evolution of B-cells into different isotype populations after the administration of three TB injections

#### Generation of Immunoglobulins (Figure 12(i))

The figure illustrates the levels of different immunoglobulins (IgM, IgG1, IgG2, and combined forms like IgG1+IgG2 and IgM+IgG) after each vaccine dose: (1) **Primary Response:** After the first dose of the vaccine, the immune system starts producing IgM as the first line of defence. IgG production begins later but is initially at lower levels; **Secondary Response:** After the second dose, there is a significant increase in both IgM and IgG levels, particularly IgG1 and IgG2. This indicates the immune system is “remembering” the antigen and mounting a stronger response; **Tertiary Response:** The third dose leads to even higher levels of IgG1 and IgG2, demonstrating a robust and heightened immune response. The rapid and strong production of immunoglobulins IgG1 and IgG2 after repeated exposure indicates that the vaccine effectively enhances immune memory and becomes more effective at recognising and neutralising the TB antigen, leading to more robust and faster antibody responses upon re-exposure. The high levels of immunoglobulins suggest that the immune system is well-prepared to clear the TB antigen quickly, reducing the risk of infection or disease progression.

#### Active B-Cell Populations (Figure 12(ii))

The figure shows the number of active B cells at different stages following each vaccine dose: **First Dose:** After the first dose, there is an increase in the population of active B cells as the immune system begins to respond to the TB antigen; **Second and Third Doses:** The population of active B cells further increases after the second and third doses, indicating that the vaccine successfully stimulates B cell activation and proliferation. This enhanced B cell activity is crucial for a strong humoral response; **Sustained Activation:** The continued presence of active B cells suggests that the immune response is maintained over time, ensuring ongoing protection against TB. The increasing number of active B cells after each vaccine dose indicates that the vaccine effectively promotes a strong humoral immune response, essential for producing antibodies that neutralise the TB pathogen. The sustained activation of B cells suggests that some cells are differentiating into memory B cells, which will provide long-term immunity by rapidly responding to future exposures to the TB antigen.

#### B Cell Isotype Evolution (Figure 12(iii))

The figure shows how B cells evolve into different isotype populations after each dose of the vaccine: **First Dose:** Initially, the majority of B cells produce IgM, which is typical of the early stages of an immune response; **Subsequent Doses:** After the second and third doses, there is a significant shift in the B cell population towards producing IgG1 and IgG2. This isotype switching reflects the maturation of the immune response and the development of a more effective and specialised antibody response; **Memory B Cells:** The presence of memory B cells is indicated by the continued production of IgG1 and IgG2 even after the antigen has been cleared, suggesting long-lasting immunity. The shift from IgM to IgG1 and IgG2 production shows that the immune system is maturing and becoming more effective at targeting and neutralising the TB pathogen. The evolution of B cells into memory B cells that produce IgG1 and IgG2 indicates that the vaccine will likely provide long-term protection by maintaining a ready population of memory B cells that can quickly respond to future infections. Thus, the TB vaccine construct effectively stimulates a robust and evolving humoral immune response. The increasing levels of immunoglobulins, the growing population of active B cells, and the isotype switching to more effective antibodies all indicate that the vaccine promotes immediate and long-term immunity.

#### Helper T Cells (HTLs) (Figure 13(i))

These are a crucial component of the immune system. HTLs assist other immune cells by secreting cytokines that enhance the ability of B cells to produce antibodies, and they help cytotoxic T cells (CTLs) to kill infected cells. Figures 13(i), 13(ii), 13(iii), and 13(iv) depict the cell-mediated immune response profiles. Figure 13 (i) shows the changes in the population of helper T cells over time, following each dose of the TB vaccine. **First Dose:** After the first exposure to the vaccine, the population of HTLs increases as the immune system begins to recognise and respond to the TB antigens; **Subsequent Doses:** With the second and third doses of the vaccine, the HTL population grows significantly. This indicates that the immune system increasingly recognises the TB antigen and is mounting a stronger response with each subsequent exposure; **Peak and Decline:** After each peak in HTL population following a vaccine dose, there is a gradual decline. This decline is expected as the immune system eliminates the antigen and reduces the active immune response. However, some HTLs persist as memory cells, crucial for long-term immunity. The increasing population of HTLs after each vaccine dose suggests that the vaccine is effectively priming the immune system to recognise and respond to TB antigens. The robust response and the generation of memory HTLs indicate that the vaccine could provide long-term protection by enabling a rapid and strong response upon re-exposure to the pathogen.

**Figure 13:**
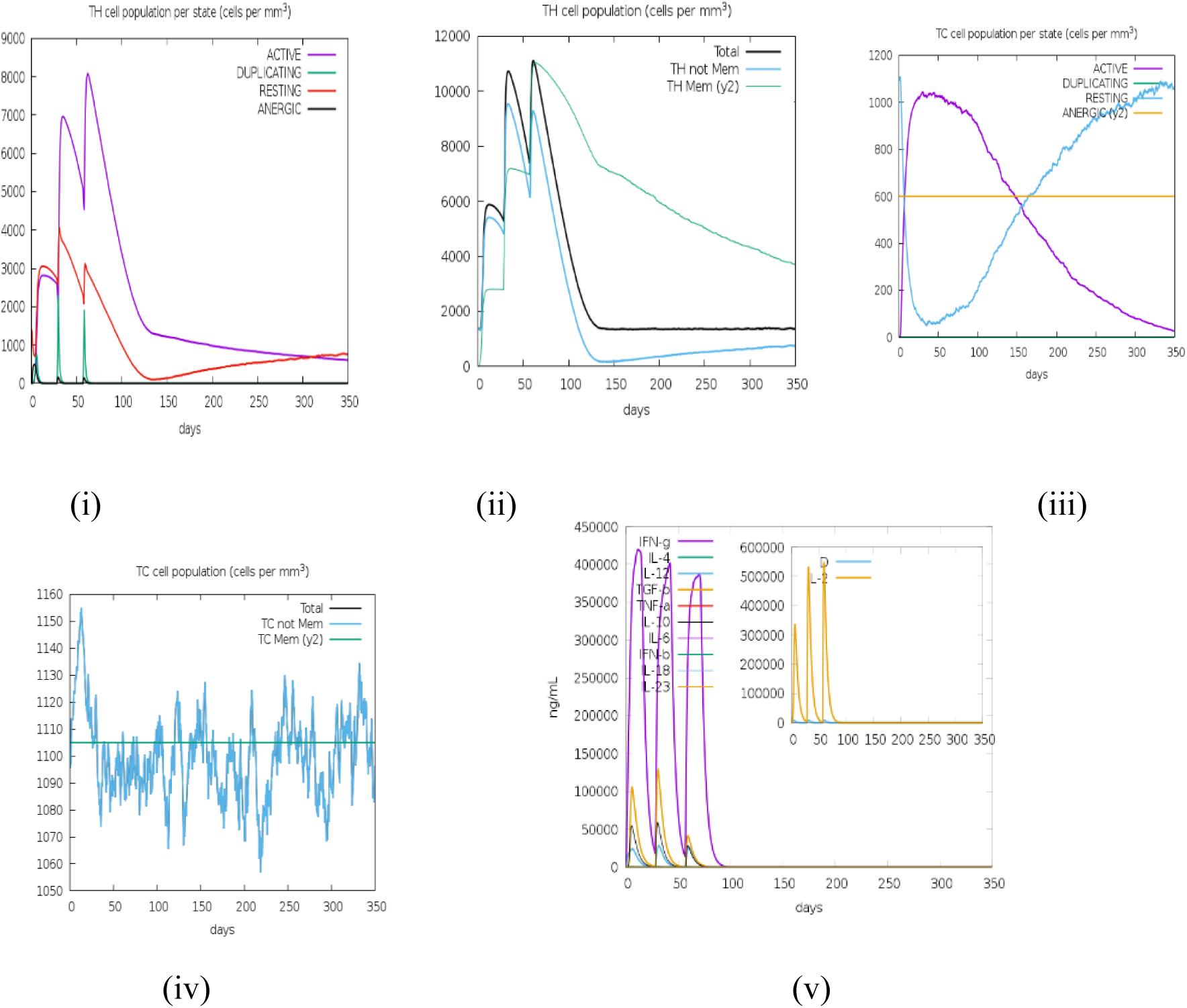
Cell-mediated immune response profile of the epitope-based TB vaccine construct: (i) population of helper T-cells per state upon exposure to TB vaccine construct, (ii) population of memory helper T-cells per state, (iii) population of cytotoxic T-cells per state after exposure to the antigen, (iv) population of memory cytotoxic T-cells per state, and (v) production of different cytokines and interleukins with Simpson index (D) (inset) in three subsequent responses.

#### Memory Helper T Cells (Figure 13(ii))

The figure shows how the population of memory HTLs evolves over time in response to the TB vaccine. **Initial Exposure:** After the first dose of the TB vaccine, the immune system generates a certain number of memory HTLs. This initial increase is due to the body’s response to the vaccine antigens; **Subsequent Exposures:** With each subsequent exposure to the vaccine (second and third doses), there is a further increase in the population of memory HTLs. This indicates that the repeated vaccine doses are successfully boosting the immune memory, leading to a stronger and more prepared immune system; **Stabilisation:** After the third dose, the memory HTL population stabilises. This stabilisation suggests that the body has developed a sufficient pool of memory cells, which will remain in the system to provide long-lasting immunity. The increasing and then stabilising population of memory HTLs shows that the vaccine is effective in creating a robust immune memory. This is important because it means that if the vaccinated individual encounters *Mycobacterium tuberculosis* in the future, their immune system will be able to respond quickly and effectively, reducing the risk of infection or severe disease.

#### Cytotoxic T Cells (Figure 13(iii))

Similar to HTLs, this figure shows the changes in the population of CTLs over time following exposure to the TB vaccine. **First Dose:** After the initial dose, there is an increase in the CTL population as the immune system starts recognizing the TB antigens; **Second and Third Doses:** The population of CTLs significantly increases after the second and third doses, indicating an enhanced immune response with repeated exposure to the antigen; **Peak and Decline:** Following each peak in CTL population, there is a decline as the immune system clears the antigen and reduces the active immune response. However, like HTLs, some CTLs differentiate into memory cells, ensuring that the body can quickly respond if the pathogen is encountered again in future. The substantial increase in CTL populations after the second and third doses suggests that the vaccine effectively stimulates a strong cell-mediated immune response, which is crucial for controlling TB infections. The generation of memory CTLs, as indicated by the persistence of CTLs even after the antigen is cleared, is essential for long-term immunity and rapid response upon re-exposure to TB.

#### Memory Cytotoxic T Cells (Figure 13(iv))

These cells are a subset of CTLs that “remember” a specific antigen (in this case, components of *Mycobacterium tuberculosis*) after the initial infection or vaccination. They can quickly respond to subsequent exposures to the same antigen by rapidly expanding and eliminating the pathogen. **Initial Response:** Upon the first exposure to the TB vaccine, the number of CTLs increases as the immune system reacts to the antigen; **Memory Formation:** After the immune response is established, some CTLs differentiate into memory CTLs; **Stabilisation:** Once the initial immune challenge has passed, the population of memory CTLs remains relatively constant. This means the body retains a pool of these cells to ensure long-term immunity. This constancy indicates successful immunisation, as it suggests that the vaccine has provoked a strong immediate immune response and established a lasting memory that will provide protection over time into the future.

The production of key cytokines and interleukins (IFN-γ, IL-4, IL-10, IL-12, IL-23, and TNF-α) was consistently high following each vaccine dose (Figure 13(v)), demonstrating effective activation of both innate and adaptive immune responses. **First Exposure:** Upon the first dose of the vaccine, there is an initial production of cytokines and interleukins, indicating the activation of the immune response; **Subsequent Exposures:** With each additional dose of the vaccine, the levels of cytokines and interleukins increase, reflecting a stronger and more sustained immune response. This repeated exposure leads to a heightened immune readiness and the ability to respond to the pathogen quickly; **IFN-γ Levels:** The high levels of IFN-γ throughout the process are significant, as they indicate the activation of macrophages and the promotion of a Th1 response, which is critical for controlling TB infections.

The Simpson Index is included in the figure (inset) to measure the diversity of the immune response. A lower value of the Simpson Index indicates greater diversity in the immune response. This is desirable because a more diverse immune response means the body can target the pathogen in multiple ways, increasing the chance of successful elimination. The figure 13(v) shows that the Simpson Index decreases over time with subsequent vaccine doses, suggesting that the immune response becomes more diverse and robust with each exposure to the vaccine.

The combined analysis of the humoral and cell-mediated immune response profiles demonstrates that the epitope-based TB vaccine construct successfully stimulates a comprehensive and robust immune response. The increase in immunoglobulins, active B cells, and T cells, along with the generation of memory cells and diverse cytokine production, suggests that the vaccine has the potential to provide long-term and effective protection against *Mycobacterium tuberculosis*. The findings from the C-ImmSim simulations highlight the vaccine’s promise in promoting both immediate and durable immunity, crucial for preventing TB infections and reducing disease burden.

## 5. Discussion and Conclusions

Tuberculosis is fast becoming incurable due to the evolution of drug-resistant bacteria and inefficacy of the currently used old vaccine leading to death of over a million affected globally. Therefore, a TB vaccine that can boost innate and humoral immunity while offering protection against drug resistant TB with universal coverage is paramount.

For decades, researchers have strived to develop an effective TB vaccine. Yet, the path to an ideal universal solution is still hindered by several challenges, such as pathogen polymorphism, eliciting a precise immune response against TB, genetic diversity of the human population and hypersensitivity. An in-depth understanding of the interaction of TB bacteria with its host, the host defence mechanism and bacterial survival strategies in evading the host immune response is important for developing an effective TB vaccine. The conventional vaccine development process is highly time-consuming, meticulous and expensive. The purification and attenuation of TB vaccine products in the laboratory is arduous and leakage of *Mycobacterium tuberculosis* is always a risk in the laboratory.

In this study, an epitope-based TB vaccine was designed by developing a conceptual framework encapsulating an in-depth understanding of host-pathogen interactions and implementing it using computational vaccinology to address the above limitations of current vaccines and challenges of the conventional vaccine development process. The potential of computational vaccinology to expedite and revolutionise the design and development of a vaccine with ease and in a short amount of time is inspiring for the field of immunology and vaccine development. Our framework utilised this potential with the following key considerations for a vaccine for global eradication of TB: (i) global coverage, fast immune response at the initial and any recurrent exposure to TB, safety for the host and maximum lethality to bacteria, maximum efficacy with minimum possible vaccine size and ease of laboratory development.

In recent years, bioinformatics tools and software have offered many significant breakthroughs in computational vaccinology. This study deployed computational vaccinology to find TB vaccine candidates with the potential to contribute significantly to the development of an effective TB vaccine with broad coverage. To achieve this, the current study focused on selecting the most highly immunodominant epitopes from *Mycobacterium tuberculosis’s* conserved and surface-exposed antigenic proteins. Various bioinformatics techniques, including comparative proteomic analysis, reverse vaccinology, immunoinformatics, and structural vaccinology, were employed to increase the likelihood of identifying the most promising vaccine targets for designing an epitope-based TB vaccine. Thus, this research offers great promise for the future of TB vaccine development.

Considering the global coverage, concern is genetic diversity of *Mycobacterium tuberculosis.* Using a single strain or single antigen does not offer a complete picture of the genetic diversity of *Mycobacterium tuberculosis*. Thus, the proteomes of all 159 completely sequenced strains of *Mycobacterium tuberculosis* available at the time were used to identify conserved proteins among those strains. Comparative proteomic analysis using a standalone BLAST identified 1982 proteins conserved across these strains, with a significant proportion (73%) involved in essential biological functions, suggesting that *Mycobacterium tuberculosis* minimizes mutations in critical proteins to maintain survival. Therefore, targeting these critical functions by a vaccine ensures both its lethality and broad spectrum coverage.

A reductionist reverse vaccinology process was performed to identify antigenic proteins within the 1982 conserved proteins. Reverse vaccinology is a cost-effective technique suitable for large-scale genomic and proteomic datasets, and it was used to scrutinize the ideal vaccine candidates by directly analysing the conserved proteins of the 159 strains of *Mycobacterium tuberculosis* with reference to H37RV strain. A total of 24 membrane-spanning, antigenic and non-allergic proteins were selected for further investigation, focussing on their potential to develop an epitope-based vaccine capable of triggering a highly specific and swift humoral and cell-mediated immune response.

The epitope-based approach provides a powerful new strategy for pathogen-specific immunity. Here, immunoinformatics approach was used to identify safe and immunodominant epitopes capable of stimulating both humoral and cell-mediated immune response in the host. From an extensive repertoire of TB epitopes predicted from immunoinformatics analysis, 27 epitopes (CTL epitopes-14, HTL epitopes-5 and B-cell epitopes-8) were shortlisted from 18 antigenic *Mycobacterium tuberculosis* proteins. These 27 epitopes were highly immunogenic, non-toxic and non-allergenic to the host. Population coverage analysis indicated that the predicted CTL and HTL epitopes could provide coverage for 99.16% of the global population, reinforcing the strategy for developing a universal TB vaccine.

Structural vaccinology then helped in developing an *in-silico* vaccine. The design of the vaccine sequence was based on a new concept introduced in this research. The docking analysis of the 27 filtered epitopes with every other epitope created 153 combinations of various epitopes. The best combination of 27 epitopes with a strong binding affinity to one another, was used for constructing vaccines. The epitopes were attached with the help of flexible linkers (GPGPG, AAY and KK). After arranging the epitopes based on their binding affinity, the final vaccine sequence was completed using two adjuvants, 50S ribosomal protein L7/L12 and β-defensin attached at the N- and C-terminals, respectively, with the help of the EAAAY linker. Post-design evaluations of antigenicity, allergenicity, and physicochemical properties confirmed the vaccine’s potential as a universal candidate satisfying the requirements for initiating an effective immune response inside the host.

The structural model of the TB vaccine was initially constructed using RaptorX and subsequently refined to enhance its quality and stability. Structural analysis confirmed that over 92% of the amino acids were positioned within the most favourable region, demonstrating high structural integrity. Further, molecular docking and dynamics simulation were performed to assess the interaction between the TB vaccine construct and TLRs (TLR-2, 4 and 6). The results indicated stable and robust binding interactions, with binding energies of -1172.8 kcal/mol (TB vaccine-TLR2), -1241.1 kcal/mol (TB vaccine-TLR4) and -1002.2 kcal/mol (TB vaccine-TLR6), suggesting the potential for initiating potent immune response. Mobility analysis further predicted minimum deformability of the vaccine construct-TLRs complexes, indicating the capacity of the constructed epitope-based TB vaccine to maintain strong interactions inside the host body, thereby effectively activating macrophages and eliciting cell-mediated and humoral immune response.

To ensure the efficient expression of the TB vaccine, codon optimisation was performed, yielding a GC content of 54.48% and a CAI of 0.99, indicating a high expression level of the designed vaccine in *E. coli*. Additionally, *in silico* cloning was done using expression vector pET-28a (+). Finally, evaluating the TB vaccine’s immune response profile provided insights into its efficacy in generating a robust and specific humoral and cell-mediated immune response. The production of B-cells, T-cells and cytokines following exposure to the TB antigen demonstrated the vaccine’s capacity to provoke a substantial immune response and foster the development of memory T-cell and B-cell populations.

The extensive analysis conducted in this study suggests that the developed epitope-based TB vaccine holds significant promise for broad-spectrum protection against diverse *Mycobacterium tuberculosis* strains. Further trials in the laboratory with *suitable in vitro* and *in vivo* models are recommended to validate the constructed TB vaccine’s safety, efficacy, and immunogenicity. The pursuit of tuberculosis eradication, alleviating the suffering of millions affected by this disease, hinges on developing such innovative vaccines. We hope this study contributes by developing a potent TB vaccine to reach this crucial goal and pave the way for a TB-free future globally.

## Supplementary Materials

